# Lowering the *HTT1a* transcript as an effective therapy for Huntington’s disease

**DOI:** 10.1101/2025.06.10.658804

**Authors:** Aikaterini Smaragdi Papadopoulou, Julia Alterman, Christian Landles, Edward J. Smith, Faith Conroy, Jemima Phillips, Maria Canibano-Pico, Iulia M. Nita, Georgina F. Osborne, Arzo Iqbal, Sarah G. Aldous, Marie K. Bondulich, Casandra Gomez-Paredes, Kirupa Sathasivam, Daniel O’Reilly, Dimas Echeverria, Konstantin Bobkov, Jonathan R. Greene, Neil Aronin, Anastasia Khvorova, Gillian P. Bates

## Abstract

Lowering the levels of *HTT* transcripts has been a major focus of therapeutic development for Huntington’s disease (HD), but which transcript should be lowered? HD is caused by a CAG repeat expansion in exon 1 of the *HTT* gene, and the rate of somatic expansion of this CAG repeat throughout life is now known to drive the age of onset and rate of disease progression. As the CAG repeat expands, the extent to which the *HTT* mRNA is alternatively processed to generate the *HTT1a* transcript and highly aggregation-prone and pathogenic HTT1a protein increases. Several HTT-lowering modalities have entered clinical trials that either target both *HTT* and *HTT1a* together, or full-length *HTT* alone. We have developed siRNAs that target the *Htt1a* mouse transcript (634/486) and used these, together with a potent *Htt*-targeting siRNA (10150) to compare the efficacy of lowering either full-length *Htt* or *Htt1a*. zQ175 and wild-type mice were treated with 10150 or 634/486 alongside control groups at 2 months of age with treatment to 6 or 10 months, or at 6 months with treatment to 10 months. The siRNA potency and durability were most effective in the hippocampus. Whilst both strategies showed benefits, despite the greater potency of 10150, targeting *Htt1a* was more effective at delaying HTT aggregation and transcriptional dysregulation than targeting full-length *Htt*. These data support HTT-lowering strategies that are designed to target the *HTT1a* transcript, either alone, or together with lowering full-length *HTT*.

**One Sentence Summary:** Lowering *HTT1a* transcript levels delays the onset of molecular and neuropathological phenotypes in a knock-in mouse model of Huntington’s disease.

## INTRODUCTION

Huntington’s disease (HD) is an inherited neurodegenerative disorder that manifests with motor abnormalities, psychiatric disturbances and cognitive decline (*1*). It is caused by a CAG repeat expansion in exon 1 of the huntingtin gene (*HTT*) that is translated to an extra-long polyglutamine (polyQ) tract in the huntingtin protein (HTT). For CAG repeats, as measured in blood, individuals with (CAG)_35_ or less will remain unaffected, (CAG)_36_ is a pathogenic repeat length and approximately (CAG)_60_ or more will result in disease onset in childhood or adolescence (*1*). CAG repeats in the mutant range expand throughout life in specific cell populations (*2, 3*), reaching several hundred CAGs in length (*3, 4*), a phenomenon known as somatic CAG repeat expansion. Neuropathologically, HTT aggregates are deposited as nuclear and cytoplasmic inclusions (*5*) and synaptic and neuronal loss occurs in the striatum, cortex and other brain regions (*6, 7*). There are no disease modifying treatments.

Although longer CAG repeats are associated with an earlier age of disease onset, genetic and environmental modifiers also contribute and several genes involved in DNA mismatch repair processes have been identified as genetic modifiers by genome-wide association studies (*8-10*). It had been known for several years that the ablation of selective mismatch repair genes in mouse models of HD prevented somatic CAG repeat expansion from occurring (*11-13*). Taken together, these data suggest that it is the rate of somatic CAG repeat expansion in neurons that drives the age of disease onset and rate of disease progression.

In the presence of an expanded CAG repeat, the *HTT* pre-mRNA can be alternatively processed through the activation of cryptic polyadenylation (polyA) sites within intron 1 (*14*). This occurs at polyA sites 2.7 kb and 7.3 kb into intron 1 for human *HTT* and 680 and 1145 base pairs into intron 1 for mouse *Htt*, generating the *HTT1a* and *Htt1a* transcripts, respectively (*14, 15*). These small transcripts encode the HTT1a protein, which has been shown, through analysis of many model systems, to be highly aggregation-prone and pathogenic (*16, 17*). The alternative processing of the *HTT* pre-mRNA, to generate *HTT1a*, increases with increasing CAG repeat length (*14, 18*), and, given this relationship, the production of the HTT1a protein may be the mechanism through which somatic CAG repeat expansion exerts its pathogenic consequences.

Targeting proximal events in the HD pathogenic cascade provides a rational therapeutic strategy and approaches to lower the levels of *HTT* transcripts are under clinical evaluation (*19*). These include agents that would lower both *HTT* and *HTT1a* e.g. AMT130, a miRNA packaged in adeno-associated virus that targets exon 1 *HTT (20, 21)*, and V0659, an antisense oligonucleotide (ASO) that targets CAG repeats (*22*). Agents that only target full-length *HTT* include the ASO tominersen (*23*) and the small molecule splicing modulators branaplam (*24*) and PTC518 (*22*). Of these, only tominersen has been evaluated in a phase III clinical trial, which failed to show any benefit (*25*).

Here, we have identified siRNAs that target mouse *Htt* intron 1 (634/486) and selectively decrease *Htt1a* levels in the brains of the zQ175 knock-in mouse model. These have been used, together with a potent siRNA that selectively decreases the full-length *Htt* transcript (10150) (*26*), to compare the effects of targeting *Htt1a* or full-length *Htt* on molecular pathogenic phenotypes. We find that, although 10150 was more potent than 634/486, targeting *Htt1a* had a more pronounced effect on delaying HTT aggregation and transcriptional dysregulation. These data support targeting *HTT1a* as a therapeutic strategy for HD.

## RESULTS

### Identification of siRNAs that target intron 1 of mouse *Htt*

We have previously published a potent siRNA that targets the 3′UTR of both human and mouse full-length huntingtin transcripts (*26*). To identify siRNAs specific to *Htt1a*, we designed and synthesized siRNAs targeting 24 sites within mouse intron 1 before the first cryptic polyA site at 680 bp (Fig 1A and table S1). The panel was initially screened for efficacy in HeLa cells expressing a luciferase reporter assay (Fig. 1B) and the 486 and 634 siRNAs, that decreased *Htt1a* to 11.6% and 8.3% of untreated levels, respectively, were selected for further analysis.

**Fig. 1.**
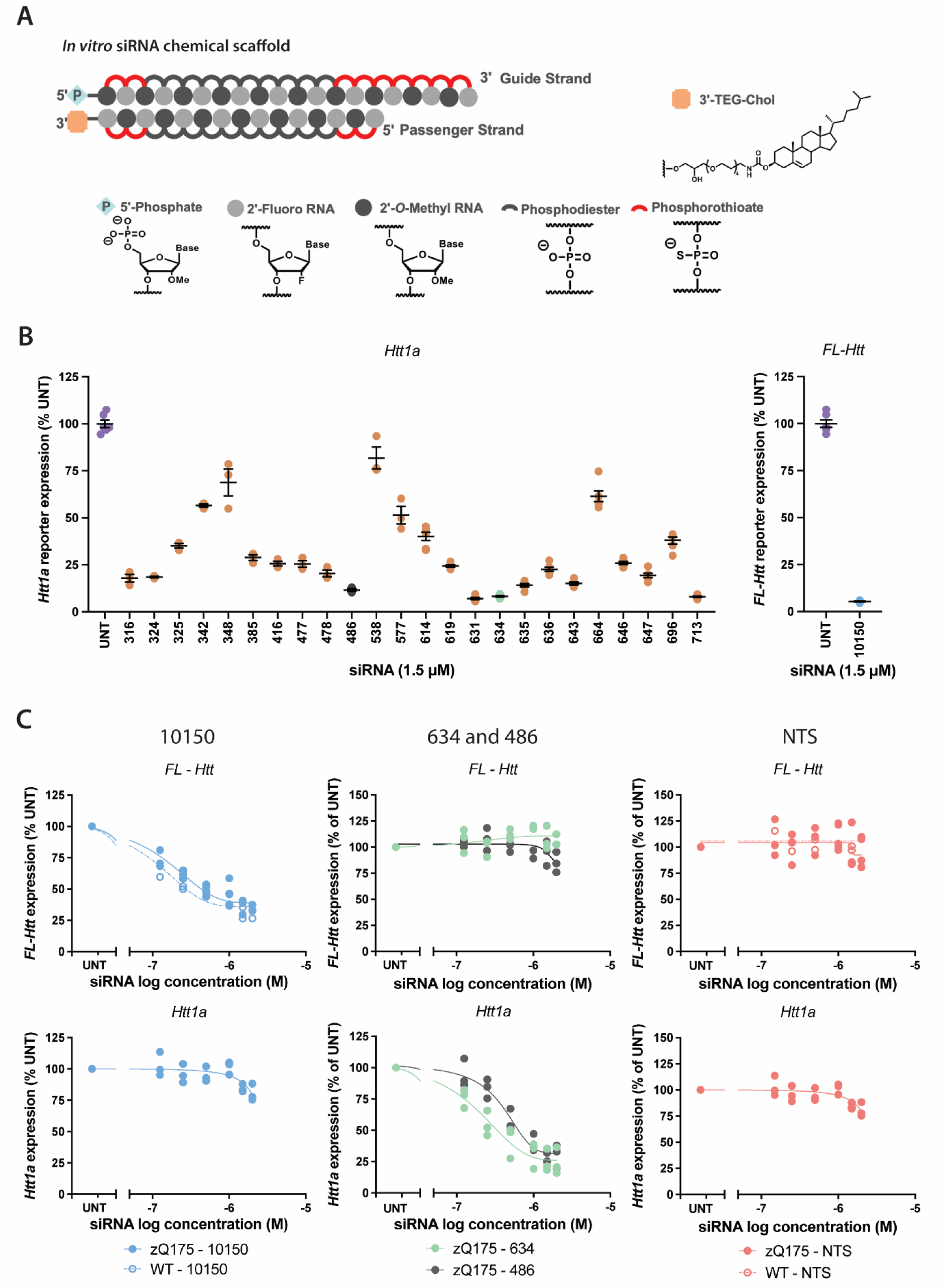
The identification of siRNAs targeting *Htt1a*. (**A**) Chemical configuration of the siRNAs used for screening. (**B**) siRNAs targeting *Htt1a* were screened in HeLa cells containing the appropriate luciferase reporter construct. The luciferase signal in *Htt1a* treated cells was compared to the untreated control. The efficiency with which the siRNA 10150 decreased full-length *Htt* levels in this reporter assay was also demonstrated. (**C**) Effect of the 10150, 634 and 486 siRNAs on the levels of *Htt* and *Htt1a* in wild-type and zQ175 MEFs as measured by the QuantiGene multiplex panel. *FL-Htt* = full-length *Htt*, NTS = non-targeting siRNA, UNT = untreated.

To determine their ability to target *Htt1a* in a more physiologically relevant system, mouse embryonic fibroblasts (MEFs), derived from wild-type and zQ175 embryos, were treated with cholesterol conjugated siRNAs: 10150, 634, 486 and a non-targeting siRNA (NTS). Except at the highest concentration, 10150 was selective for full-length *Htt* and 634 and 486 were selective for *Htt1a*, with the 634 siRNA being the most potent. The NTS had no effect (Fig. 1C).

### The 10150 siRNA and 643/486 siRNAs are specific for their targets *in vivo*

We have previously described a divalent siRNA modification that supports potent, sustained gene silencing in the central nervous system (CNS) of mice and non-human primates following a single injection into the cerebral spinal fluid (CSF) (*27*). Therefore, divalently modified versions of 10150 targeting full-length *Htt*, 634 and 486 targeting *Htt1a* as well as an NTS were synthesized for *in vivo* studies (Fig. 2A and table S2).

**Fig. 2.**
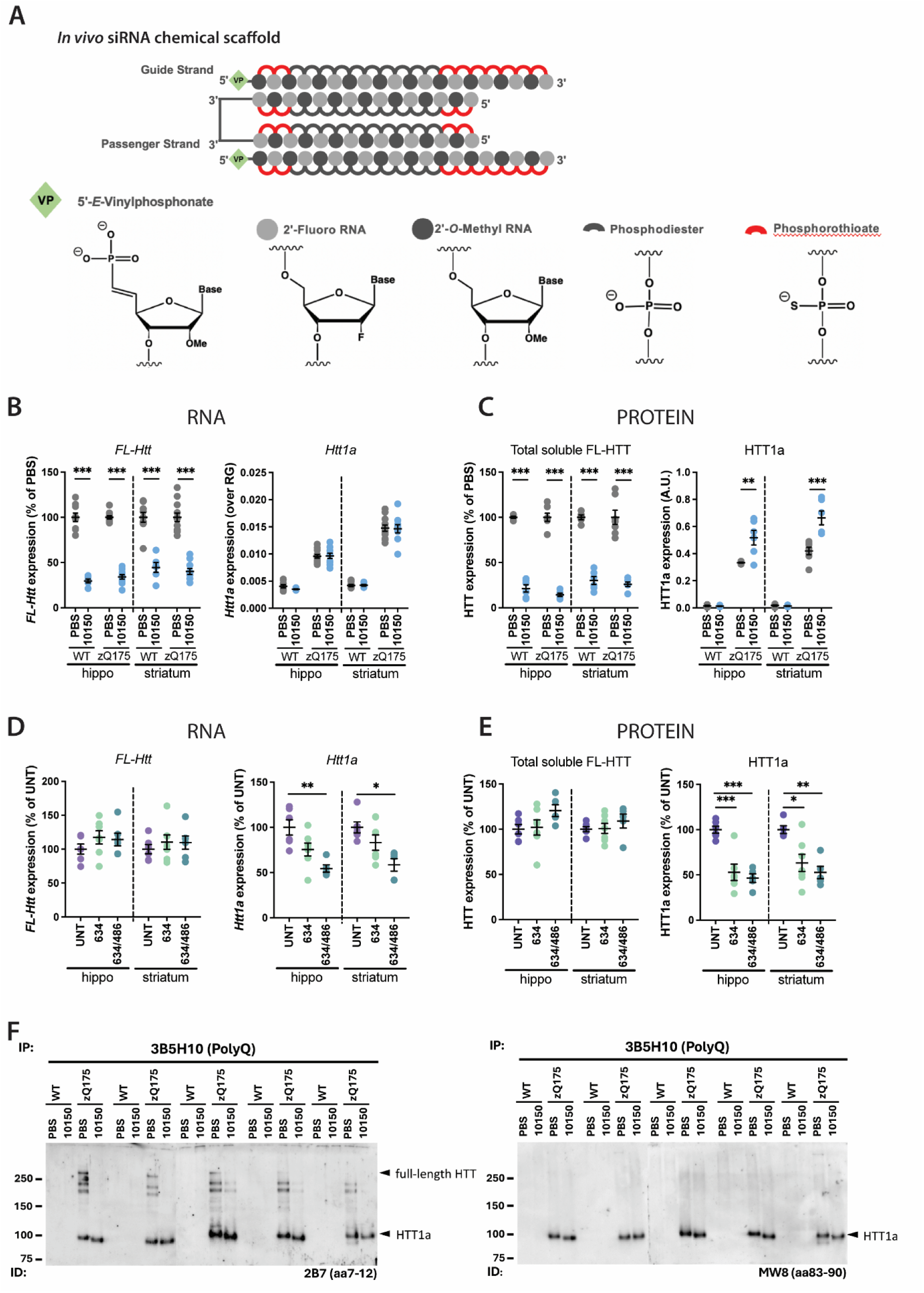
Selective reduction of full-length *Htt* by 10150 and *Htt1a* by 634/486 *in vivo*. (**A**) Chemical configuration of the divalent siRNAs used for the *in vivo* trials. (**B**) Full-length *Htt* transcripts were decreased in the hippocampus and striatum in response to 10150 treatment but *Htt1a* levels were unaltered (n = 6-12/ treatment, 2-7/ gender/ treatment). (**C**) HTRF analysis demonstrated that 10150 treatment decreased full-length HTT levels (MAB5490-MAB2166) and appeared to increase HTT1a (2B7-MW8) (n = 5-6/ treatment, 2-4/ gender/ treatment). (**D**) Treatment with 634/486 decreased *Htt1a* but not full-length *Htt* levels in the hippocampus and striatum (n = 3-4/ gender/ treatment). **(E**) HTRF analysis demonstrated that 634 and 634/486 decreased HTT1a protein levels (2B7-MW8) and not full-length HTT (MAB5490-MAB2166) (n = 3-4/ gender/ treatment). (**F**) Immunoprecipitation of mutant HTT with the 3B5H10 antibody from wild-type and zQ175 cortical lysates, treated with PBS or 10150, and immunodetected with 2B7 showed that full-length HTT and its proteolytic fragments had been depleted by treatment with 10150. Immunodetection with MW8, that is specific for HTT1a, showed that the levels of HTT1a were unaltered by 10150 treatment (n = 5/ treatment/ genotype). Protein size standards in kilodaltons. Statistical analysis was one-way ANOVA with Tukey’s post hoc correction or Kruskal-Wallis with Dunn’s post hoc correction. Error bars: mean ± SEM. \**P* ≤ 0.05, \*\**P* ≤ 0.01, \*\*\**P* ≤ 0.001. aa = amino acids, FL = full-length, Hippo = hippocampus, NTS = non-targeting siRNA, UNT = untreated, WT = wild type.

Wild-type and zQ175 mice were treated with 10150 to ensure that the expected reduction in the level of full-length *Htt* was achieved. Wild-type and zQ175 mice were injected bilaterally into the lateral ventricles (ICV) with 20 nmole (10 nmole / ventricle) 10150 or with phosphate buffer saline (PBS) at 8 weeks of age and sacrificed at 12 weeks. A QuantiGene multiplex panel (*28*) was used to assess the level of full-length *Htt* and *Htt1a* mRNAs in brain regions. Full-length *Htt* was decreased to comparable levels in wild-type and zQ175 animals and to a greater extent in the hippocampus as compared to the striatum, cortex, and cerebellum (Fig. 2B and fig. S1A). Reductions in full-length HTT protein were assessed by homogeneous time resolved fluorescence (HTRF) and found to be ∼86% in the hippocampus, 74% in the striatum, 69% in the cortex and 45% in the cerebellum of zQ175 mice (Fig. 2C and fig. S1B). The decrease in HTT protein was slightly greater than the reduction in mRNA levels, consistent with previous observations (*29*). Treatment with 10150 had no effect on *Htt1a* mRNA levels (Fig. 2B and fig S1A), but, surprisingly, the reduction in full-length HTT appeared to have resulted in an increase in HTT1a protein in all four brain regions (Fig.2C and fig. S1B).

The *Htt1a* transcript is not generated in wild-type mice (*14, 18*), and therefore, the potency of siRNAs 634 and 486 was only assessed in zQ175 animals. Given that the 634 siRNA had been more potent than 486 *in vitro*, we opted to test 634 alone (10 nmole / ventricle) and the 634 and 486 siRNAs in combination (5 nmole of each / ventricle) as compared to untreated animals. zQ175 mice were treated by bilateral ICV injection at 8 weeks of age and sacrificed at 12 weeks as before. The use of the siRNAs in combination proved to be more potent than 634 alone (Fig. 2D and fig. S1C), with the reduction in HTT1a protein being 54% in the hippocampus, 47% in the striatum, 47% in the cortex and 28% in the cerebellum (Fig. 2E and fig. S1D). Treatment with 634/486 had no effect on full-length huntingtin mRNA or protein levels (Fig. 2D and E and fig. S1C and D).

### Treatment with 10150 does not lead to an increase in HTT1a levels

To explore the apparent increase in HTT1a protein with the reduction of full-length HTT, we performed immunoprecipitation and western blotting as an independent assessment of the effect of the 10150 siRNA on HTT and HTT1a protein levels. Under these denaturing conditions, treatment with 10150 decreased the level of full-length HTT and its proteolytic fragments but HTT1a levels remained the same and were not increased (Fig. 2F). We considered that the apparent increase in HTT1a could be an artefact of the HTRF assay due to limiting antibody concentrations; upon the removal of full-length HTT, there may have been more antibody available to detect HTT1a. To explore this hypothesis, we immunodepleted full-length HTT by immunoprecipitation with either the MAB5490 or MAB2166 antibody from either wild-type or zQ175 lysates and measured full-length HTT and HTT1a levels by HTRF (fig. S2). This confirmed that full-length HTT levels had been depleted (fig. S2B), however, HTT1a levels were unchanged (fig. S2C). Therefore, the apparent increase in HTT1a in response to HTT depletion upon 10150 treatment was not an artefact of the HTRF assay. We do not yet understand the mechanism for the apparent increase in HTT1a, but do not think that it reflects total HTT1a levels.

### An interventional trial to assess the efficacy of targeting *Htt* or *Htt1a*

Our *in vivo* pilot data indicated that treatment with 10150 decreased full-length HTT levels by ∼70-85% in hippocampus, striatum and cortex. The siRNAs targeting *Htt1a* were less potent, but the combination of 634 and 486 had resulted in a reduction of HTT1a of ∼50% in these forebrain regions. Therefore, we designed an interventional study to compare the effects of depleting full-length *Htt* or *Htt1a* (Fig. 3). Treatment was initiated at 2 months of age, prior to the onset of transcriptional and aggregation phenotypes, or at 6 months of age once these phenotypes are well-established (*30, 31*). There were four treatment groups per cohort: untreated, NTS, 10150 (full-length *Htt*) or the 643/486 combination (*Htt1a*) (total of 10 nmole / ventricle). Three cohorts consisted of: treatment at 2 months and sacrificed at 6 months (EARLY); at 2 months, again at 6 months and sacrificed at 10 months (DOUBLE); and at 6 months and sacrificed at 10 months (LATE) (n = 18/gender/treatment group/genotype) (Fig. 3).

**Fig. 3.**
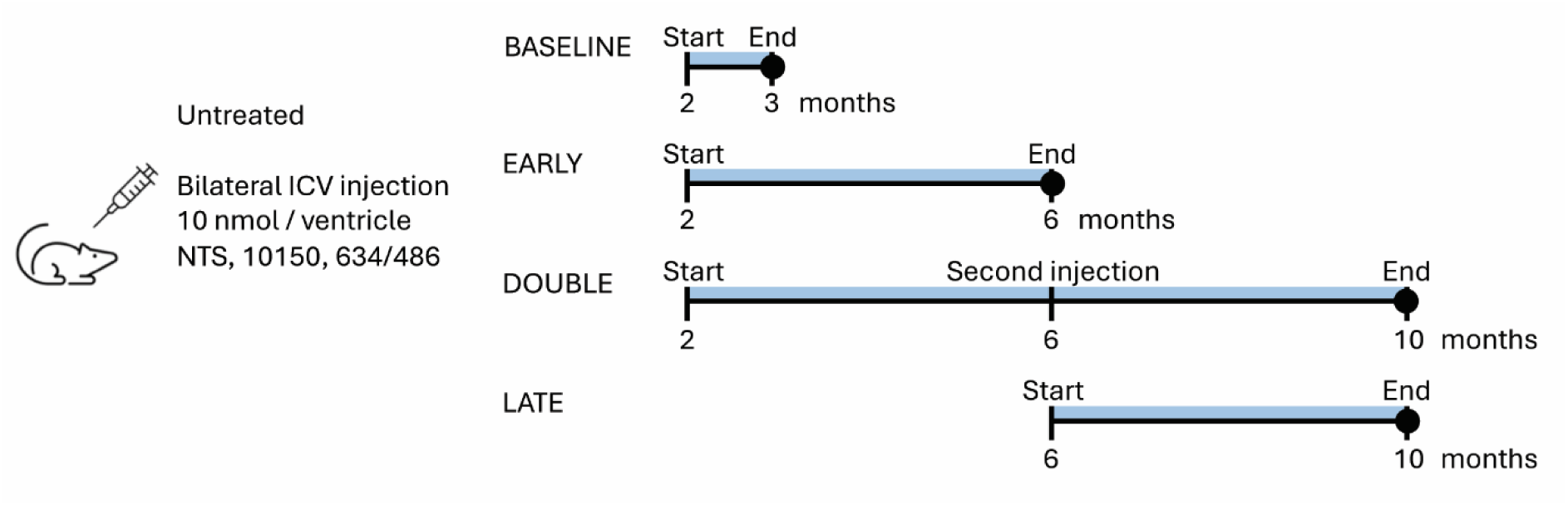
Design of the interventional trial to compare the efficacy of lowering full-length *Htt* or *Htt1a*. Wild-type and zQ175 mice were randomly sorted into four treatment groups: untreated, NTS, 10150 and 634/486. Bilateral ICV administration was with 10 nmole divalent siRNA / ventricle. Mice were treated at 2 months of age and sacrificed at 3 months, to measure the extent to which full-length *Htt* and *Htt1a* levels had been decreased at baseline. To determine the effects of full-length *Htt* or *Htt1a* reduction on molecular phenotypes, mice were treated at 2 months of age and sacrificed at 6 months (EARLY cohort), at 2 months, again at 6 months and sacrificed at 10 months (DOUBLE cohort) and at 6 months and sacrificed at 10 months (LATE cohort). NTS = non-targeting siRNA.

A cohort of wild-type and zQ175 mice was also treated at 2 months of age and sacrificed at 3 months (BASELINE) (Fig. 3), to assess the potency of the siRNAs that had been synthesized for the interventional study (n = 7/gender/treatment group/genotype). Treatment with 10150 (fig. S3A and B) resulted in a reduction in full-length HTT protein of ∼80% in the hippocampus, 56% in the striatum and 58% in the cortex of zQ175 mice (fig. S3B). Treatment with 634/486 led to a reduction in the HTT1a protein of 51% in the hippocampus, 24% in the striatum and 26% in the cortex (fig. S3D). As before, the siRNAs had the greatest effect in the hippocampus and the 634/486 siRNA combination was less potent than 10150.

### Sustained reduction of *Htt* and *Htt1a* was most pronounced in the hippocampus

We began by assessing the extent to which the huntingtin transcripts and HTT protein isoforms were decreased, four months after the siRNA dose administered at 2 or 6 months of age in the EARLY, DOUBLE or LATE cohorts. The durability of the siRNA treatment was greatest in the hippocampus (Fig. 4). The NTS had no effect on *Htt* or *Htt1a* transcript levels or on HTT or HTT1a protein levels respectively for any of the three treatment regimens (Fig. 4). Treatment with 10150 resulted in sustained reduction in the full-length *Htt* transcript (Fig. 4A), as well as the full-length HTT protein (Fig. 4B). The decrease in HTT protein in wild-type and zQ175 mice was 38% and 30% in the EARLY, 65% and 60% in the DOUBLE and 42% and 62% in the LATE cohorts, respectively. The mean extent of knock-down was less in the EARLY cohort and the wild-type LATE cohort mice because of the considerable variability among animals. Treatment with 634/486 (Fig. 4C and D) led to a sustained reduction in HTT1a protein in the EARLY (37%), DOUBLE (45%) and LATE (41%) cohorts (Fig. 4D).

**Fig. 4.**
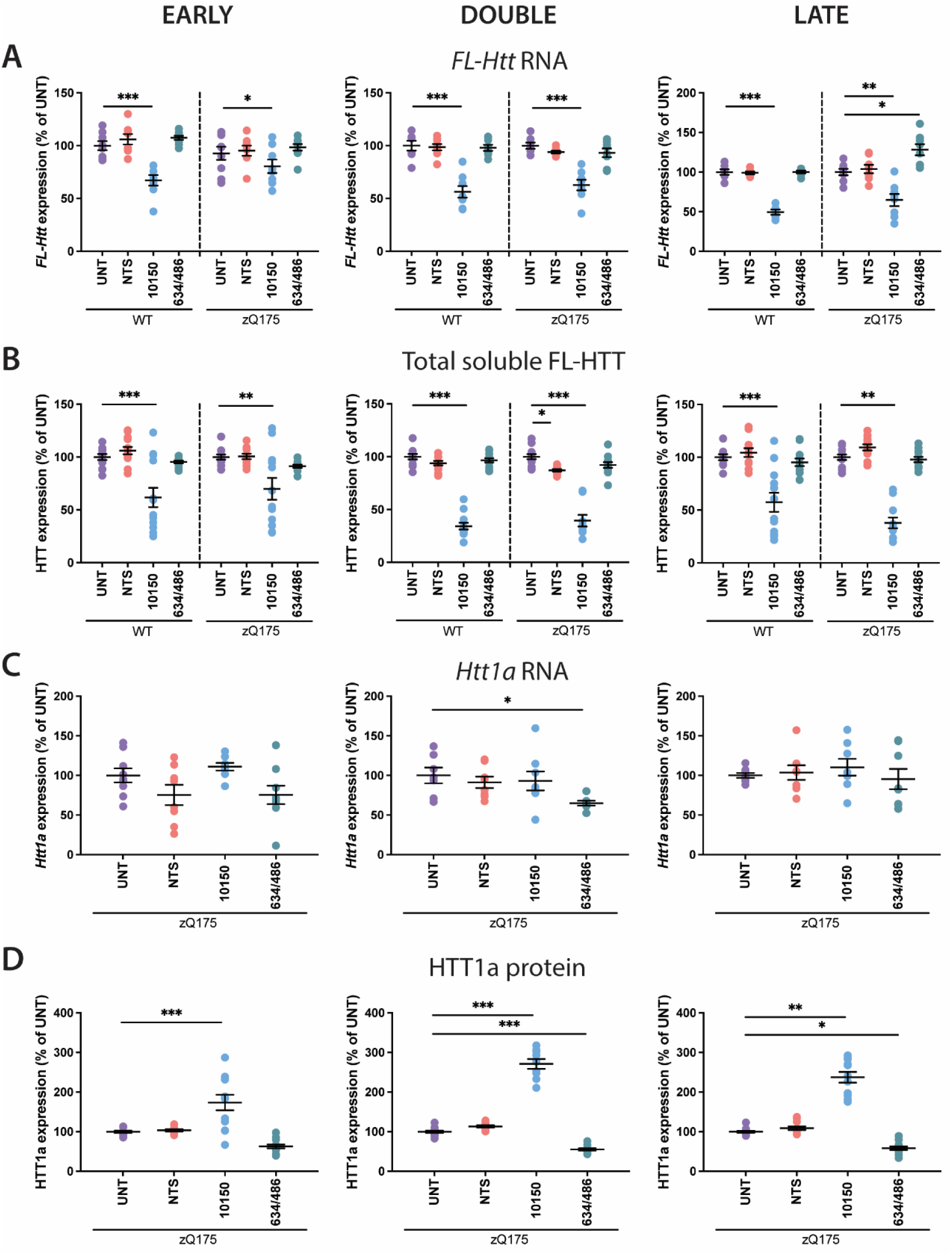
Effect of treatment with 10150 or 634/486 on huntingtin mRNA and protein levels in the hippocampus four months post-injection. Wild-type and zQ175 mice, either untreated, or 4 months post-treatment with NTS, 10150 or 634/486 in the EARLY, DOUBLE or LATE cohorts. (**A**) Full-length *Htt* mRNA levels as measured by the ‘short 3’UTR’ probe on the QuantiGene multiplex panel. (**B**) Full-length HTT protein levels as measured by the HTRF MAB5490-MAB2166 assay. (**C**) *Htt1a* mRNA levels as measured by the I_1_-pA_1_ probe on the QuantiGene multiplex panel. (**D**) HTT1a protein levels as measured by the HTRF 2B7-MW8 assay. (**A-D**) RNA and protein analyses were performed with separate sets of mice: RNA, n = 3-5/ gender /treatment; protein, n = 5-7 /gender /treatment. Statistical analysis was one-way ANOVA with Tukey’s post hoc correction or Kruskal-Wallis with Dunn’s post hoc correction. Error bars: mean ± SEM. \**P* ≤ 0.05, \*\**P* ≤ 0.01, \*\*\**P* ≤ 0.001. FL = full-length, NTS = non-targeting siRNA, WT = wild type UNT = untreated.

A decrease in full-length HTT levels 4 months after treatment with 10150 was also maintained in the striatum (fig. S4) and cortex (fig. S5), but to a lesser extent than in the hippocampus. Treatment with 634/486 did not result in a sustained reduction in HTT1a levels in the striatum (fig. S4) and in the EARLY cohort in the cortex (fig. S5).

### Targeting *Htt1a* delayed nuclear HTT aggregation in the hippocampus

The potency and durability of the 10150 and 634/486 siRNAs had proved greatest in the hippocampus, and therefore we prioritized this brain region to investigate the treatment effects on histological and molecular phenotypes. HTT aggregation can be detected in 10 CNS regions of the zQ175 mouse brain by 1 month of age and, as such, is the earliest phenotype to develop in this mouse model (*31*). It is apparent by immunohistochemistry before 2 months of age in the striatum and CA1 region of the hippocampus, and by 3 months of age in the dentate gyrus (*31*). Hippocampal sections from all four treatment groups from each of the EARLY, DOUBLE and LATE cohorts were immunostained with the S830 anti-HTT antibody. Treatment with the NTS had no effect on hippocampal HTT aggregation in any of the three cohorts (Fig. 5).

**Fig. 5.**
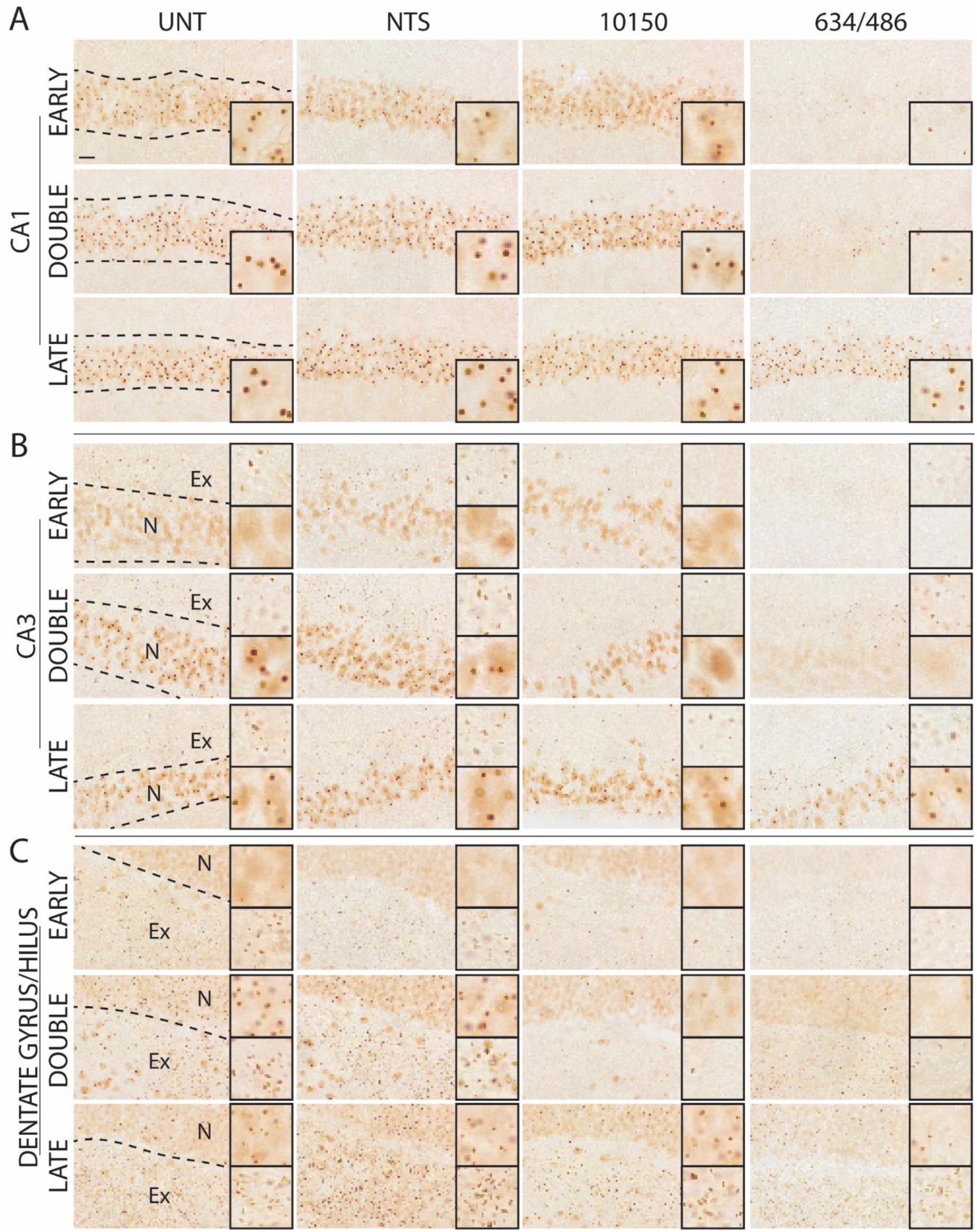
Targeting *Htt* or *Htt1a* has differential effects on HTT aggregation in the hippocampus. Coronal hippocampal sections immunostained with S830 and imaged at the level of the (**A**) CA1, (**B**) CA3 and (**C**) dentate gyrus / hilus from the EARLY, DOUBLE and LATE cohorts (n = 3/ treatment). N = nuclear, Ex = extranuclear. In zoomed images, nuclear aggregation is on bottom in (B) and top in (C) and neuropil inclusions are on top in (B) and bottom in (C. Scale bar = 20 µm, zoomed images are 20 µm^2^. NTS = non targeting siRNA, UNT = untreated.

The CA1 pyramidal cells of the untreated sections contained diffuse HTT aggregation and an inclusion body at 6 months of age, and by 10 months, the inclusion body was larger and more pronounced, with a concomitant reduction in diffuse aggregation (Fig. 5A). The decrease in full-length HTT with 10150 treatment had no effect on HTT aggregation in the CA1 region in any of the three cohorts (Fig. 5A). In contrast, reduction in HTT1a, through 634/486 treatment at 2 months of age led to a dramatic reduction in HTT aggregation in the EARLY and DOUBLE cohorts (Fig. 5A).

HTT aggregation in the CA3 pyramidal cell nuclei of untreated sections was predominantly diffuse with few nuclear inclusions at 6 months, and by 10 months, this had progressed such that all nuclei contained an inclusion body (Fig. 5B). The reduction in HTT1a levels through treatment with 634/486 almost prevented the appearance of the diffuse aggregation at 6 months in the EARLY cohort and by 10 months of age, had dramatically decreased the level of diffuse aggregate staining and prevented the appearance of nuclear inclusions in the DOUBLE cohort (Fig.5B). Treatment with 10150 at 2 months of age had decreased the appearance of nuclear inclusions at 10 months in the DOUBLE cohort. Treatment with either 634/486 or 10150 had also decreased the concentration of extranuclear inclusions, lying dorsal to the nuclear layer at 6 and 10 months of age (Fig. 5B).

In the untreated sections, HTT aggregation appeared diffuse in the dentate gyrus nuclei at 6 months of age, progressing to include nuclear inclusions by 10 months (Fig. 5C). A dense concentration of neuropil aggregation in the hilus was already apparent by 6 months of age, which increased over the next 4 months (Fig. 5C). Reduction of HTT1a by 634/486 treatment at 2 months of age prevented the formation of nuclear aggregation in the dentate gyrus at 6 months and confined this to faint nuclear stain with no inclusion bodies at 10 months (Fig. 5C). Treatment at 2 months of age with 634/486 or 10150 almost ablated the presence of extranuclear inclusions in the hilus at both 6 and 10 months of age (Fig. 5C).

Treatment at 6 months of age with 10150, or with 634/486 appeared to have a much more modest effect on HTT aggregation in the CA1 (Fig. 5A), CA3 (Fig. 5B) or dentate gyrus and hilus (Fig.5C).

### The effects of targeting full-length *Htt* or *Htt1a* on HTT aggregation in the hippocampus are complex

To define the effects of treatment with 10150 or 634/486 on hippocampal HTT aggregation in more detail, we employed an unbiased quantification of nuclear inclusions, diffuse nuclear aggregation and extranuclear inclusions (Fig. 6). We applied this to the DOUBLE cohort, as by 10 months of age, nuclear inclusions had formed in all three regions, and treatment from 2 months of age with either 10150 or 634/486 appeared to have had pronounced effects (Fig. 5). In the nuclei of the CA1 pyramidal cells, decreasing HTT1a with 634/486 almost prevented the formation of diffuse aggregation and nuclear inclusions, whereas decreasing full-length HTT had no discernable effect (Fig. 6A). In the neuronal nuclei of the CA3 and the dentate gyrus, both treatment with 10150 or 634/486 substantially reduced the formation of nuclear inclusions, but only 634/486 also decreased the level of diffuse nuclear aggregation (Fig. 6A). Both treatment with 10150 or 634/486 considerably decreased the number of extranuclear inclusions in the CA3 and the hilus (Fig. 6A).

**Fig. 6.**
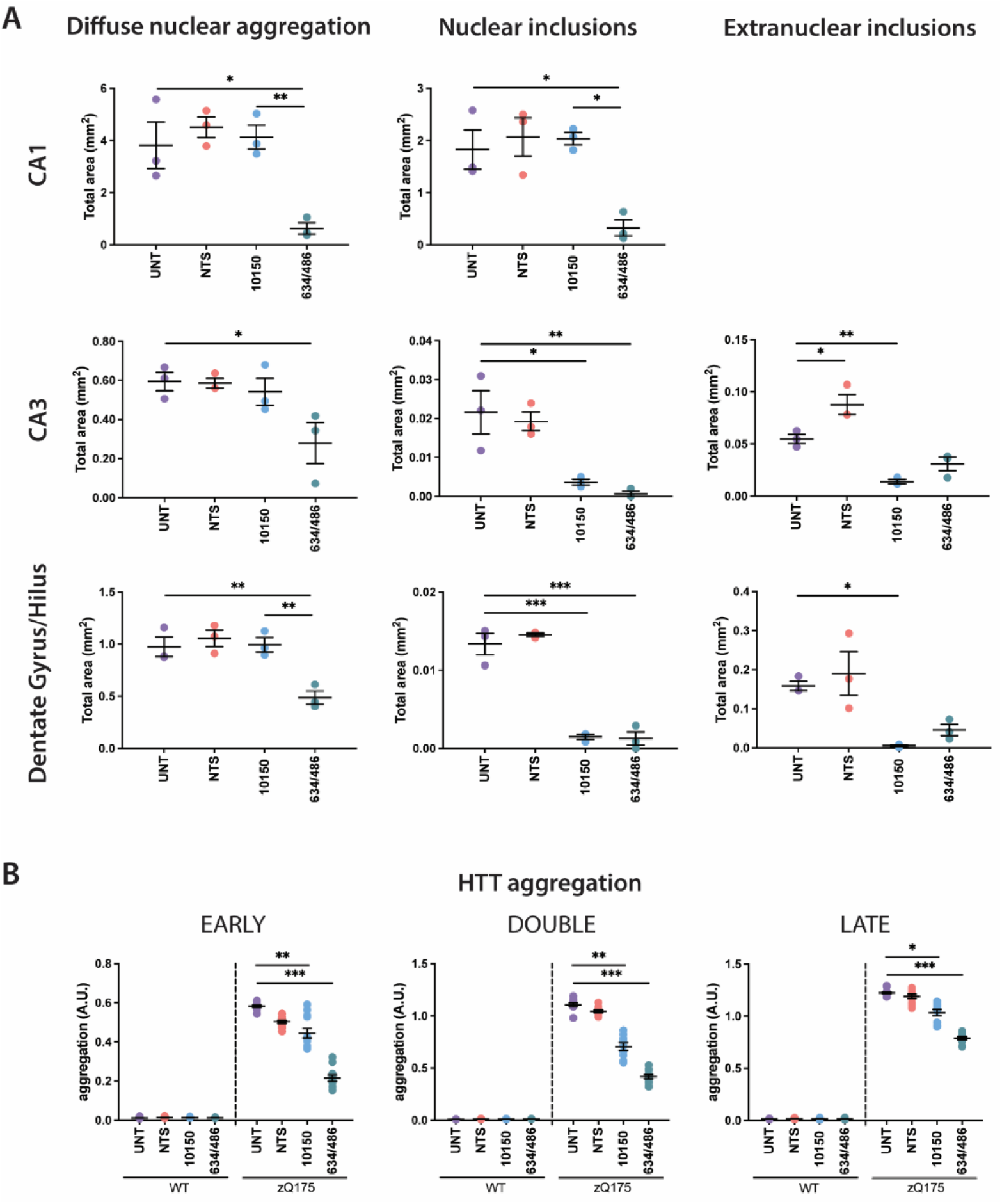
Lowering *Htt1a* has a more pronounced effect on HTT aggregation in the hippocampus than lowering full-length *Htt*. (**A**) Quantification of the total area nuclear inclusions, diffuse nuclear aggregation and extranuclear inclusions in images acquired from the CA1, CA3 and dentate gyrus for the four treatment groups of the DOUBLE cohort (n = 3/ treatment). (**B**) HTT aggregation as measured by the HTRF 4C9-MW8 assay (n = 12/ treatment). Statistical analysis was by one way ANOVA with Tukey’s post hoc correction. Error bars are mean ± SEM. \**P* ≤ 0.05, \*\**P* ≤ 0.01, \*\*\**P* ≤ 0.001. AU = arbitrary units, NTS = non-targeting siRNA, UNT = untreated.

We used HTRF (antibody pair 4C9-MW8) to quantify HTT aggregation levels at the whole tissue level. Consistent with the immunohistochemical data, treatment with 10150 or 634/486 led to a considerable reduction in HTT aggregation in the EARLY and DOUBLE cohorts with 634/486 having the most pronounced effect (Fig. 6B). HTRF analysis confirmed that the reduction in HTT aggregation in the LATE cohort, after initiation of treatment at 6 months of age was more modest than starting at 2 months.

### Targeting *Htt1a* delayed transcriptional dysregulation in the hippocampus more than targeting *Htt*

As transcriptional dysregulation is a consequence of aggregated HTT in the nucleus (*32, 33*), we sought to determine whether treatment with siRNAs had partially restored the wild-type transcriptome. Transcriptional dysregulation in the hippocampus is less well defined than in the striatum but had previously been reported for zQ175 mice at 6 and 10 months of age (*30*). Hippocampal RNA was sequenced, and we began by comparing the NTS and untreated transcriptomes for zQ175 and wild-type mice for each of the EARLY, DOUBLE and LATE cohorts. The NTS had no effect on the transcriptional profile for either wild-type or zQ175 mice in the EARLY or LATE cohorts (fig. S6). For the DOUBLE cohort, the expression of 73 genes was altered in NTS-treated wild-type mice, and 47 in NTS-treated zQ175 mice (fig. S6 and S7). Therefore, the NTS data sets for the DOUBLE cohort were included in further analyses.

Comparison of the transcriptional profile of untreated zQ175 or NTS-treated zQ175 mice with their wild-type counterparts was highly correlated with previously published data (Fig. 7A). Analysis of full-length *Htt* and *Htt1a* levels confirmed that these transcripts were decreased 4 months post-dosing with 10150 or 634/486, respectively, in all three cohorts (Fig. 7B), although there was a considerable variability between animals in some cases. Treatment at 2 months of age with either 10150 or 634/486 led to an improvement in the dysregulated profile in the EARLY and DOUBLE cohorts, although the effect was more pronounced with the reduction of *Htt1a* than with that of full-length *Htt* (Fig. 7C). Treatment at 6 months of age led to little improvement, and in the case of either 10150 or 634/486 was as likely to have led to an exacerbation of the level of dysregulation (Fig. 7C).

**Fig. 7.**
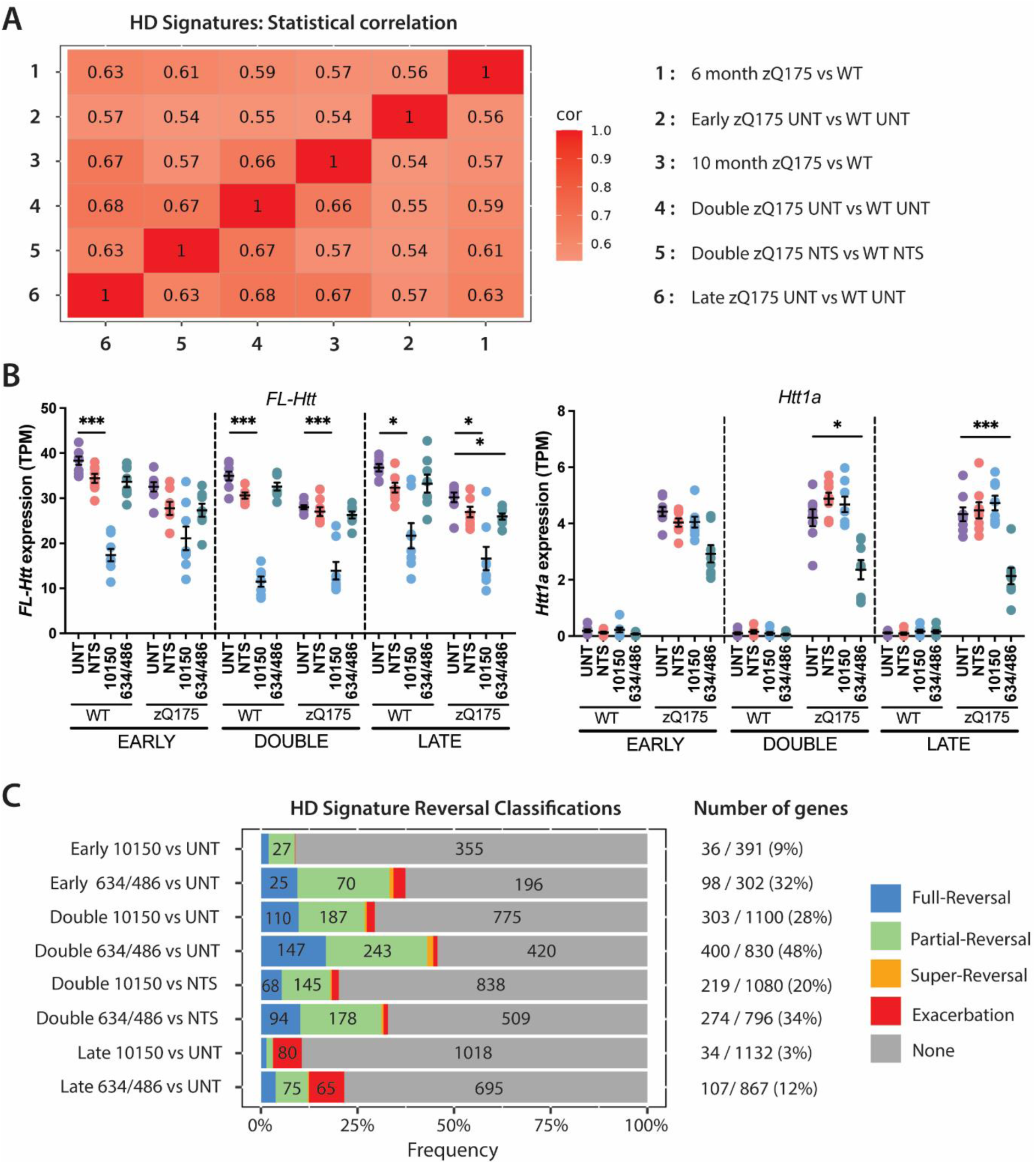
Lowering *Htt1a* delays transcriptional dysregulation in the hippocampus to a greater extent than lowering full-length *Htt*. (**A**) Heatmap showing that the dysregulated hippocampal transcriptional profile in the EARLY, DOUBLE and LATE cohorts (represented by the DESeq2 Wald statistic) is highly correlated to previously published comparisons of transcriptional profiles between zQ175 and wild-type mice at 6 and 10 months of age (*30*). (**B**) The extent to which the reduction in full-length *Htt* and *Htt1a* has been sustained four months post-dosing with 10150 or 634/486 in the hippocampal RNA-seq datasets from the EARLY, DOUBLE and LATE cohorts. (**C**) The proportion of dysregulated genes for which the level of expression was reversed toward wild-type levels for mice treated with either 10150 or 634/486 as compared to untreated mice for each of the cohorts and compared to NTS treated mice for the DOUBLE cohort. N = 8/ treatment. Statistical analysis was by DESeq2 with Benjamini-Hochberg correction. Error bars are mean ± SEM. \**P* ≤ 0.05, \*\*\**P* ≤ 0.001. Cor = correlation, FL = full-length, NTS = non-targeting siRNA, TPM = transcripts per million, UNT = untreated, WT = wild type.

### Reduction in full-length HTT levels had no effect on HTT aggregation or transcriptional dysregulation in the striatum

Treatment with 10150 led to a reduction in full-length HTT protein in the striatum of ∼56%, 1-month post-dosing (fig.S3A), with the level of knock-down remaining at ∼30% at 10 months of age in the DOUBLE cohort (fig. S4A). As this reduction in HTT might be comparable to that achieved in a clinical trial, we investigated whether it might have improved striatal HTT aggregation or transcriptional dysregulation phenotypes. In contrast, the level of HTT1a reduction in the striatum was only 23% 1-month post-dosing with 634/486 with no sustained reduction in any of the cohorts (fig. S4B).

Coronal striatal sections from untreated mice or those treated with the NTS or 10150 in the EARLY, DOUBLE or LATE cohorts were immunoprobed with the S830 antibody. At 6 months of age, aggregation in the zQ175 striatum includes diffuse nuclear aggregation, nuclear inclusions and extranuclear inclusions (*31*) and by 10 months, the level of nuclear diffuse staining has decreased, and the inclusion size increased. Treatment with 10150 had no effect on HTT aggregation in any of the three cohorts (Fig 8A), which was confirmed with the HTRF HTT aggregation assay (4C9-MW8) (Fig. 8B).

**Fig. 8.**
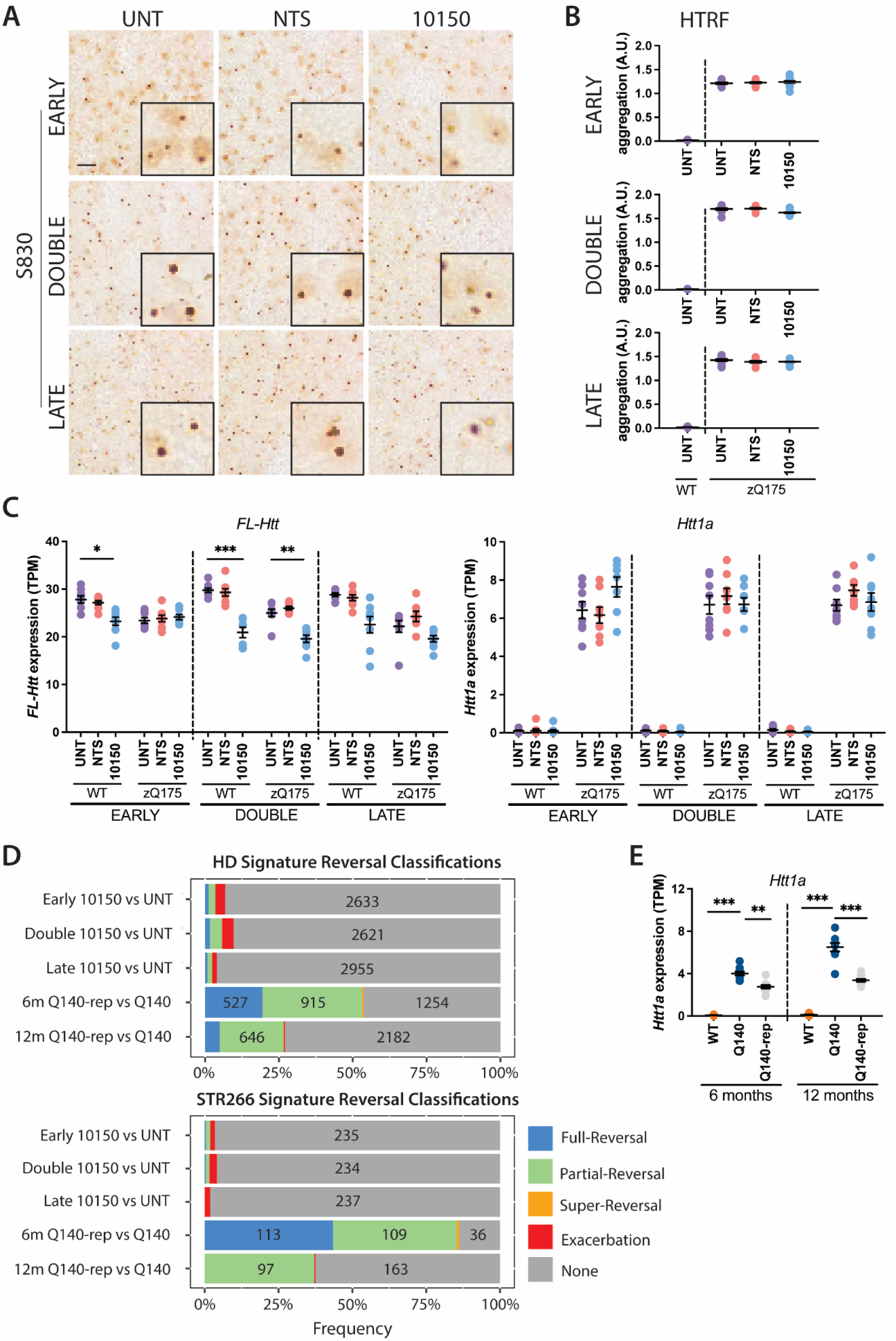
Reduction in full-length HTT had no effect on HTT aggregation or transcriptional profiles in the striatum. (**A**) Coronal striatal sections immunostained with S830 from the EARLY, DOUBLE and LATE cohorts (n = 3/treatment). Scale bar = 20 µm, zoomed images are 20 µm^2^. (**B**) HTT aggregation as measured by the HTRF 4C9-MW8 assay (n = 12/ treatment). (**C**) At 4 months post-dosing, there is a sustained reduction in full-length *Htt*, but not *Htt1a* in the striatum of mice from the DOUBLE cohort treated with 10150 in the RNA-seq data sets (n = 8/ treatment). (**D**) Reversal of transcriptional dysregulation profile for all dysregulated genes (top panel) and for the striatal signature of 266 genes (bottom panel). The reduction of full-length *Htt* in the zQ175 mice treated with 10150, had no effect on the dysregulated profiles. A decrease in the expression of the mutant *Htt* gene by ∼50% in Q140 mice through the Lac repressor from 2- to 6- and 2- to 12-months of age resulted in a profound reversal of these dysregulated profiles. (**E**) The Lac repression led to a decrease in *Htt1a* by ∼50% in the Q140 mice (n = 9-10/ genotype). Statistical analysis was by one way ANOVA with Tukey’s post hoc correction or by or by DESeq2 with Benjamini-Hochberg correction. Error bars are mean ± SEM. \**P* ≤ 0.05, \*\**P* ≤ 0.01, \*\*\**P* ≤ 0.001. M = months, NTS = non-targeting siRNA, Rep = repressed, TPM = transcripts per million, UNT = untreated.

Dysregulation of the striatal transcriptome has been well-documented, with the expression level of several thousand genes altered in the zQ175 striatum at 6 and 10 months of age (*30*). Striatal RNA was sequenced for untreated animals and those treated with NTS and 10150 from the three cohorts (n = 8 / treatment group). Treatment with the NTS had no impact on the zQ175 or wild-type transcriptome for all three cohorts (fig. S8) and comparison of the striatal transcriptional profile in untreated zQ175 as compared to untreated wild-type mice was consistent with that observed in previously published data sets (fig. S9).

Analysis of the RNA-seq datasets confirmed that a sustained decrease in full-length *Htt* could be detected in zQ175 mice in the DOUBLE cohort at the end of the trial, and that *Htt1a* levels were unchanged (Fig. 8C). There were 2,895 dysregulated genes in the zQ175 striatum of untreated mice in the DOUBLE cohort at 10 months of age. Although, treatment with 10150 reversed the expression of 6% of these toward wild-type levels, for 4% of genes the dysregulation was exacerbated (Fig. 8D). The effect of 10150 treatment was similar when considering the ‘striatal signature’ comprising 266 robustly dysregulated genes across multiple HD models (*34*), with 2% reversed toward wild-type levels and 2% exacerbated. The effects of decreasing *Htt* and *Htt1a* have previously been studied in Q140 knock-in mice that express the *lac* repressor and in which a *lac* operator has been placed upstream of the *Htt* promoter on the mutant allele (*35*). We analyzed the historical RNA-seq data sets for mice in which mutant *Htt* gene expression in Q140 striatum was repressed from 2 to 6, and from 2 to 12 months of age by ∼50% (*35*). Whilst the reduction in full-length *Htt* was comparable to treatment with 10150, the *lac* repressor also led to a reduction in *Htt1a* levels (Fig. 8E). At 6 months of age the expression of >50% of dysregulated genes were reversed toward wild-type levels, comprising >85% of ‘striatal signature’ genes (Fig. 8D). Therefore, the improvements reported with the *lac* repressor were predominantly achieved through the reduction in HTT1a and not full-length HTT.

## DISCUSSION

Somatic CAG repeat expansion drives the age of onset and rate of disease progression for HD and is the first step in the pathogenic cascade. The generation of the *HTT1a* transcript presents a likely candidate for the next step, as *HTT1a* levels increase with increasing CAG repeat length and this transcript encodes the highly pathogenic HTT1a protein. We have previously shown that preventing somatic expansion in zQ175 mice, with repeat expansions of ∼185 CAGs, had no effect on the striatal HTT aggregation or transcriptional dysregulation (*36*). Therefore, as expansion beyond (CAG)_185_ did not exacerbate these phenotypes, zQ175 mice provide an excellent knock-in model for the preclinical testing of treatments targeting the next step in the pathogenic cascade. Here we have used siRNAs to compare the effects of lowering either the full-length HTT or HTT1a proteins in zQ175 mice. We found that, although less potent, siRNAs targeting *Htt1a* were more effective at delaying HTT aggregation and transcriptional dysregulation phenotypes than an siRNA targeting full-length *Htt*.

The greatest effects of 10150 or 634/486 on HTT and HTT1a levels, respectively, occurred in the hippocampus. Treatment with 10150 led to a considerable reduction in the full-length HTT protein whereas the reduction in HTT1a was never greater than 50%. By immunohistochemistry, nuclear aggregation is first detected as a diffuse immunostain throughout the nucleus, which may progress to forming a nuclear inclusion. Treatment with 634/486 from 2 months of age greatly delayed HTT aggregation in the neuronal nuclei of the CA1, CA3 and dentate gyrus. Even at 10 months of age, after 8 months of treatment, the level of diffuse aggregation and the presence of nuclear inclusions was extremely low in all three hippocampal regions. These results are in keeping with a model in which HTT1a nucleates HTT aggregation in nuclei (*31*). Treatment with 10150 had no effect on nuclear inclusions in the CA1, but interestingly, almost prevented their formation in the CA3 and dentate gyrus, whilst at the same time, the level of diffuse aggregation remained unchanged. Therefore, at least in the CA3 and dentate gyrus, full-length HTT is contributing to the aggregation process in nuclei, and its depletion has slowed the trajectory through diffuse aggregation to the formation of inclusions. The effect of 10150 or 634/486 on transcriptional dysregulation was consistent with the extent to which HTT aggregation in hippocampal nuclei was delayed; treatment with 634/486 delayed the extent to which transcription was dysregulated in zQ175 mice more effectively than 10150, despite the lower potency. Treatment at 6 months of age, once HTT aggregation is well-established, was much less effective and the impact on transcriptional dysregulation was very small.

Treatment with either 10150 or 634/486 also resulted in a profound reduction in the number of neuropil aggregates in the CA3 and hilus. In the hilus, these inclusions will have originated from cell bodies in the dentate gyrus, and therefore, the formation of both nuclear and extranuclear inclusions was delayed by targeting full-length HTT or HTT1a in this brain region.

These data indicate that the interplay between full-length HTT and HTT1a protein levels in the initiation of HTT aggregation is complex and differs between specific types of neurons. As the CAG repeat increases through somatic expansion, the levels of the HTT1a protein increase, whilst the level of full-length HTT decreases (and consequentially, the levels of proteolytic HTT fragments) (*18*). Therefore, the relative concentration of HTT1a and small proteolytic fragments of HTT is dynamic within a neuron and changes as the CAG repeat expands. HTT aggregation is a concentration-dependent process for which the longer the polyQ length, the lower the threshold concentration required for nucleation (*37*), and for a given polyQ length, HTT1a (the exon 1 HTT protein) aggregates faster than other N-terminal HTT fragments (*17*). We have previously shown that HTT1a and small proteolytic fragments of HTT are lost from the soluble phase of brain lysates as HTT aggregation proceeds (*38*). We speculate that both HTT1a and small proteolytic fragments of HTT contribute to HTT aggregation and that full-length HTT levels must be lowered to a greater extent than HTT1a to impact the aggregation process.

We have shown that decreasing HTT1a levels is more effective at delaying molecular phenotypes in an HD mouse model than decreasing full-length HTT. These results complement those in a companion study demonstrating that ASOs that lower both *Htt1a* and full-length *Htt* prevented HTT aggregation and transcriptional dysregulation in the *Hdh*Q111 mouse models of HD, whereas those targeting full-length *Htt* alone had no effect (Bragg et al. this issue). Together, these data suggest that it was predominantly the reduction of *Htt1a* and not full-length *Htt* that was responsible for the benefits to a wide range of molecular, physiological and behavioral phenotypes when the expression of the *Htt* gene was decreased through a *lac* repressor strategy (*35*) or by the AAV delivery of zinc finger proteins coupled to a transcriptional repressor (*39*). To date, there has been one phase III clinical trial of an ASO targeting full-length *HTT*, which was terminated prematurely because individuals receiving ASO treatment were not meeting their primary endpoints, and potentially performing worse than those on placebo (*25*). Our data do not support HTT-lowering strategies that target full-length HTT alone. Instead, they support treatments that will lower both HTT and HTT1a, as demonstrated by miRNA (*20, 21*), ZFPs (*40*) or ASOs (Bragg et al. this issue), or the development of new agents that selectively target human HTT1a.

## MATERIALS AND METHODS

### Study Design

The objective of the study was to assess the effects of lowering the full-length *Htt* or *Htt1a* transcripts on molecular and immunohistochemical phenotypes in the zQ175 mouse model of HD. The experimental unit was either a wild-type or zQ175 mouse embryonic fibroblast line (MEF), or a wild-type or zQ175 mouse. The experimental design is summarized in Fig. 3. Sufficient siRNA was synthesized to ensure that the baseline study and each of the treatment cohorts could be performed using the same batch of drug. Wild-type and zQ175 mice were imported from the CHDI colonies at the Jackson laboratory at 7 weeks of age in groups of 48 or 64 animals per week. Each week, mice were sorted randomly into equal numbers of males and females for the four cohorts: untreated, NTS, 10150 and 634/486. In total, there were 36 mice / genotype mice in each treatment group to ensure that there were sufficient animals dedicated to each of the molecular and histological assessments. The number of mice used for QuantiGene analysis (*28*), HTRF (*18, 41*) and RNA-seq (*30*) as well as immunohistochemistry (*31*) were based on previous experimental data. All data were collected blind to genotype and treatment. Any outliers were determined statistically.

### Mouse breeding and maintenance

All procedures were performed in accordance with the Animals (Scientific Procedures) Act, 1986 and approved by the University College London Ethical Review Process Committee. For the isolation of MEFs, zQ175 males were backcrossed to C57BL/6J females (Charles River, Netherlands). For siRNA treatment, heterozygous zQ175 mice and wild-type littermates were imported from the CHDI Foundation colonies at the Jackson Laboratory (Bar Harbor, Maine) at 7 weeks of age. Mouse husbandry and health monitoring were as previously described (*42*). Animals were kept in individually ventilated cages with Aspen Chips 4 Premium bedding (Datesand) and environmental enrichment comprising chew sticks and a play tunnel (Datesand). All mice had constant access to water and food (Teklad global 18% protein diet, Envigo, the Netherlands). Temperature was regulated at 21°C ± 1°C and mice were kept on a 12 h light/dark cycle. The facility was barrier-maintained and quarterly non-sacrificial FELASA (Federation of European Laboratory Animal Science Associations) screens found no evidence of pathogens.

### Genotyping and CAG repeat sizing

Tail biopsies were collected upon sacrifice for genotype confirmation and genotyped as previously described (*41*). CAG repeat sizing had been performed by the CHDI Foundation before shipment to UCL and CAG repeat lengths are summarized in table S3.

### Oligonucleotide Synthesis

Oligonucleotides were synthesized on a Dr. Oligo 48 (Biolytic) for 1-μmol *in vitro* scale or a MerMade12 (Biosearch Technologies) for *in vivo* (10-20) µmol scale using standard solid-phase phosphoramidites (*43*). Phosphoramidites used include: 2’-Fluoro RNA or 2’-*O*-Methyl RNA modified phosphoramidites (ChemGenes and Hongene Biotech), 5’-(*E*)-Vinyl tetra phosphonate (pivaloyloxymethyl) (Hongene), and 2’-O-methyl-uridine 3’-CE phosphoramidite (Hongene). Divalent oligonucleotides (DIO) were synthesized on a custom solid support prepared in-house (*27*) Oligonucleotide synthesis and deprotection was performed as previously described (*44*).

### Cell Culture for oligonucleotide screening

Oligonucleotides were initially screened against a renilla luciferase reporter assay in HeLa cells as previously described (*44*). Cells were plated at 2 million cells per 10 cm^2^ dish 18 h prior to transfection with 20 mg reporter plasmid and 60 mL Lipofectamine 2000 (Invitrogen) in serum-free OptiMEM (Gibco) for 6 h at 37°C. Cells were rinsed with PBS and incubated overnight in DMEM (Thermo Fisher) with 10% fetal bovine serum (FBS; Gibco). After 18 h, cells were transferred to a 96 well plate and treated with 1.5 µM cholesterol conjugated siRNA for 72 h. Cells were washed twice with PBS and lysed with Dual-Glo reagent for 15 min. Dual-Glo Stop & Glo reagent was added and after 15 min, luminescence was read on a luminometer. Luminescence values were normalized to untreated controls.

For follow-up screening, mouse embryonic fibroblasts (MEFs) were isolated from E14.5 zQ175 and wild-type embryos and maintained in DMEM supplemented with 10% FBS and 1% penicillin-streptomycin in a humidified incubator at 37°C with 5% CO^2^. MEFs were transformed with the viral oncogene Simian Virus-40 large tumor antigen (SV40 T-antigen) using the Cell Immortalization Kit (CILV01 Alstem) according to manufacturer’s recommendations and as previously described (*42*). Three zQ175 MEF lines (Z3, Z4 and Z7), that had CAG repeat lengths of 191, 196 and 194, respectively, and two wild-type cell clones were chosen for downstream applications. Wild-type and zQ175 MEFs were seeded at 25,000 cells per well in 96-well plates in DMEM with 6% FBS (Gibco) and 1% penicillin and streptomycin (Gibco). After 72 h, siRNAs were added in a 1:1 ratio of DMEM and OptiMEM (Gibco) with 3% FBS. 72 h later, cells were lysed for QuantiGene according to the manufacturer’s protocol.

### Surgery

Mice were deeply anaesthetized with isoflurane and bilaterally microinjected by stereotaxic injection with 20 nmole siRNA per mouse in 5 µL per lateral ventricle at 500 nL per minute. The coordinates from bregma were: -0.2 mm AP, ± 0.8 mm ML, -2.5 mm DV. Mice were observed for 10 days post-surgery and bi-weekly thereafter. For molecular analyses, tissues were harvested, snap frozen in liquid nitrogen and stored at -80°C.

### QuantiGene Analysis

The lysis of tissue samples and the QuantiGene multiplex assays were carried out as described (*28*). Data analysis was performed by normalizing raw MFI signals obtained to the geometric mean of the housekeeping gene signals. The results of the ‘short 3′UTR’ and ‘I_1_-pA_1_’ probes for full-length *Htt* and *Htt1a*, respectively, were graphed for illustrative purposes.

### Antibodies

The antibodies used for HTRF, immunoprecipitation, western blotting and immunohistochemistry are summarized in table S4.

### Homogeneous time-resolved fluorescence (HTRF)

HTRF assays were performed as previously described (*31, 41*). Lysate dilutions and antibody concentrations are summarized in table S5.

### Western blotting and immunoprecipitation

Immunoprecipitations from wild-type and zQ175 cortical samples were performed using the anti-polyglutamine 3B5H10 antibody (Sigma-Aldrich) as described previously (*38*). For western blotting, proteins were denatured, separated by 7.5% SDS-Polyacrylamide Acrylamide Gel Electrophoresis (SDS-PAGE), blotted onto nitrocellulose membranes, and detected by chemiluminescence, as described previously (*33, 38*). Primary antibody dilutions were 1:1000.

### Immunohistochemistry

Mice were transcardially perfusion fixed with 4% paraformaldehyde (Pioneer Research Chemical Ltd) and tissue processing and the storage of sections was as previously documented (*33*). The immunostaining and imaging of coronal sections using S830 was carried out as previously outlined except the ABC reagent was diluted 1:4 (*31*). Sections from all cohorts were stained, imaged and processed together. To perform an analysis of aggregation throughout the hippocampus, three images were captured across four hemispheres for the CA1 and the dentate gyrus, and a single image from four hemispheres for the CA3 and the hilus. Images were acquired as previously described (*36*). For quantification and thresholding, images were converted to 8-bit in ImageJ and a signal intensity level was chosen to indicate staining above background. Signals were segmented based on pixel intensity and the number of positive pixels in contact with each other. A gate of 2 pixels was applied to all analyses. For nuclear inclusions, objects were considered positive for all signal over 190 for the CA1 and 150 for the CA3 and dentate gyrus. For diffuse nuclear aggregation, positive objects were signals over 210 – 190 for the CA1 and 210 – 150 for the CA3 and the dentate gyrus. For extranuclear inclusions positive signals were more than 205.

### RNA-sequencing

RNA was prepared and sequenced as previously described (*36*). Differential expression tests were performed in R using DESeq2 (*45*) also as previously described (*36*); a posterior probabilities-based method (*35*) was used to quantify the probability that the siRNA treatments restored transcriptional dysregulation, either partially or fully, on a gene-by-gene basis. The ‘DOUBLE NTS versus Untreated’ dysregulated genes lists was tested for ranked gene set enrichment against the GOBP (gene ontology biological process) (*46*) version 18.9.29 using the GSEA function within the R clusterProfiler (*47*) package. The DESeq2 Wald statistic was used to rank genes and a clusterProfiler *q* < 0.05 was deemed significant. siRNA guide sequences nucleotides 2-8 (designated mer7m8) were used to scan 3’UTRs throughout the transcriptome for potential off-target effects using the R SeedMatchR package (*48*). Genes containing > 0 mer7m8 matches (data file S1) were excluded from the analysis of restored transcriptional dysregulation described above.

### Statistics

Data were screened for outliers at sample level using the ROUT test (Q = 1%; GraphPad Prism v9) and outliers were removed from the analysis. All data sets were tested for a normal gaussian distribution (Shapiro-Wilk, Prism v9). Statistical analysis was by one-way ANOVA with either Tukey’s *post hoc* tests. Non-parametric analysis was performed by the Kruskal-Wallis test with Dunn’s *post hoc* analysis. Graphs were prepared using Prism v9 (GraphPad Software, California, USA). P-values ≤ 0.05 were considered statistically significant.

## List of Supplementary Materials

Fig. S1. Selective reduction of full-length *Htt* by 10150 *in vivo*.

Fig. S2. Immunodepletion of HTT from cortical lysates does not result in an increase in HTT1a.

Fig. S3. Effect of treatment with 10150 or 634/486 on huntingtin mRNA and protein levels one-month post-injection.

Fig. S4. Effect of treatment with 10150 or 634/486 on HTT or HTT1a levels in the striatum four months post-injection.

Fig. S5. Effect of treatment with 10150 or 634/486 on HTT or HTT1a levels in the cortex four months post-injection.

Fig. S6. Hippocampal transcriptional response to treatment with the non-targeting siRNA.

Fig. S7. Hippocampal transcriptional response to treatment with the non-targeting siRNA in the DOUBLE cohort.

Fig. S8. Striatal transcriptional response to treatment with the non-targeting siRNA.

Fig S9. Striatal transcriptional profile is comparable to that in previously published data sets.

Table S1. Sequence and modifications of oligonucleotides used for the luciferase screen.

Table S2. Sequence and modifications of oligonucleotides used for *in vivo* experiments.

Table S3. CAG repeat length for each of the treatment groups.

Table S4. Antibodies.

Table S5. Antibody and lysate concentrations for HTRF assays.

Data file S1. List of off-target seed-matches for the siRNAs.

References (49-57) (numbers for references only cited in Supplementary Material).

## Acknowledgments

We thank Sarah Allen for collating the sequence information for the siRNAs and Sam Hildebrand for advice on the analysis off-target effects of siRNAs. We thank Brenda Lager and Britt Callahan for providing the zQ175 and wild-type mice from the CHDI Foundation colony at the Jackson Laboratory and Viktoria Andreeva for the GEO submission.

## Funding

This work was funded by:

CHDI Foundation A-11004 (GPB)

CHDI Foundation A-5038 (NA)

CHDI Foundation A1781 (NA)

Wellcome Trust 223082/Z/21/Z (GPB)

National Institutes of Health S10 OD036329 (AK)

National Institutes of Neurodegenerative Disease and Stroke NS104022-06 (AK)

National Institutes of Health U01 NS114098 (NA)

Hereditary Disease Foundation Postdoctoral Fellowship (DO)

## Author contributions

Conceptualization: AK, GPB, ASP, JRG.

Methodology: JFA, ASP, CL, EJS.

Investigation: ASP, JFA, FC, CL, EJS, GFO, JP, MC-P, IMN, AI, SGA, MKB, CG-P, KB, DO, KS, DME.

Visualization: ASP, CL, EJS.

Funding acquisition: AK, GPB, NA.

Project administration: AK, NA, GPB, JRG.

Supervision: AK, GPB, NA, ASP, CL, EJS, JRG.

Writing – original draft: GPB, ASP, CL, EJS.

Writing – review & editing: all authors

## Data and materials availability

RNA-seq data that support the findings of this study have been deposited in GEO with the accession number GSE284504. The authors confirm that all other data supporting the findings in this study are available within the article and its Supplementary Material and Data File. Raw data will be shared by the corresponding author upon request.

