## Supplemental Material for "Lowering the *HTT1a* transcript as an effective therapy for Huntington’s disease"

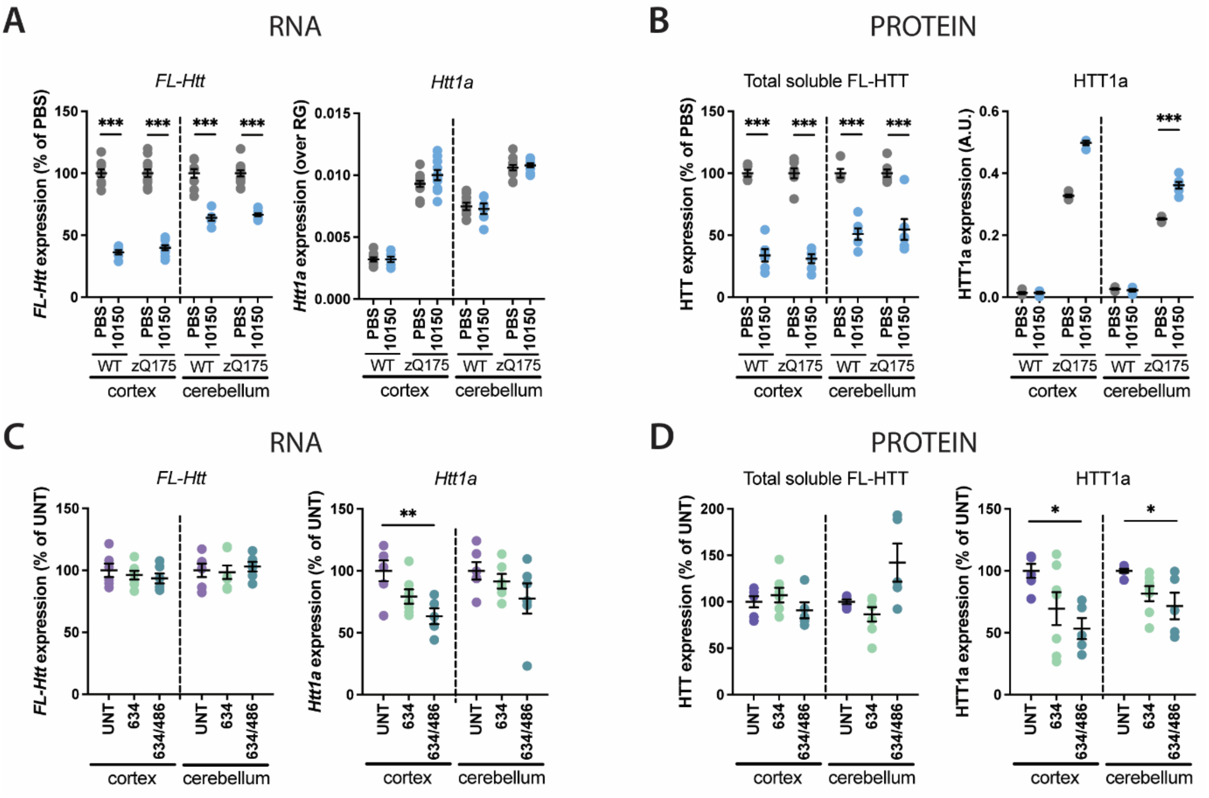


**Fig. S1 Selective reduction of full-length *Htt* by 10150 and *Htt1a* by 634/486 *in vivo*.** (**A**) Full-length *Htt* transcripts were decreased in the cortex and cerebellum in response to 10150 treatment but *Htt1a* levels were unaltered (n = 6-12/ treatment, 2-7/ gender/ treatment). (**B**) HTRF analysis (MAB5490-MAB2166) demonstrated that 10150 treatment decreased full-length HTT levels and appeared to increase HTT1a (n = 5-6/ treatment, 2-4/ gender/ treatment). (**C**) Treatment with 634/486 decreased *Htt1a* but not full-length *Htt* levels in the cortex and had no effect on *Htt1a* in the cerebellum (n = 3-4/ gender/ treatment). **(D**) 634/486 decreased HTT1a protein levels and not full-length HTT (n = 3-4/ gender /treatment). Statistical analysis was one-way ANOVA with Tukey's post hoc correction. Error bars: mean ± SEM. $^{**}P \leq 0.01$, $^{***}P \leq 0.001$. FL = full-length, WT = wild type.

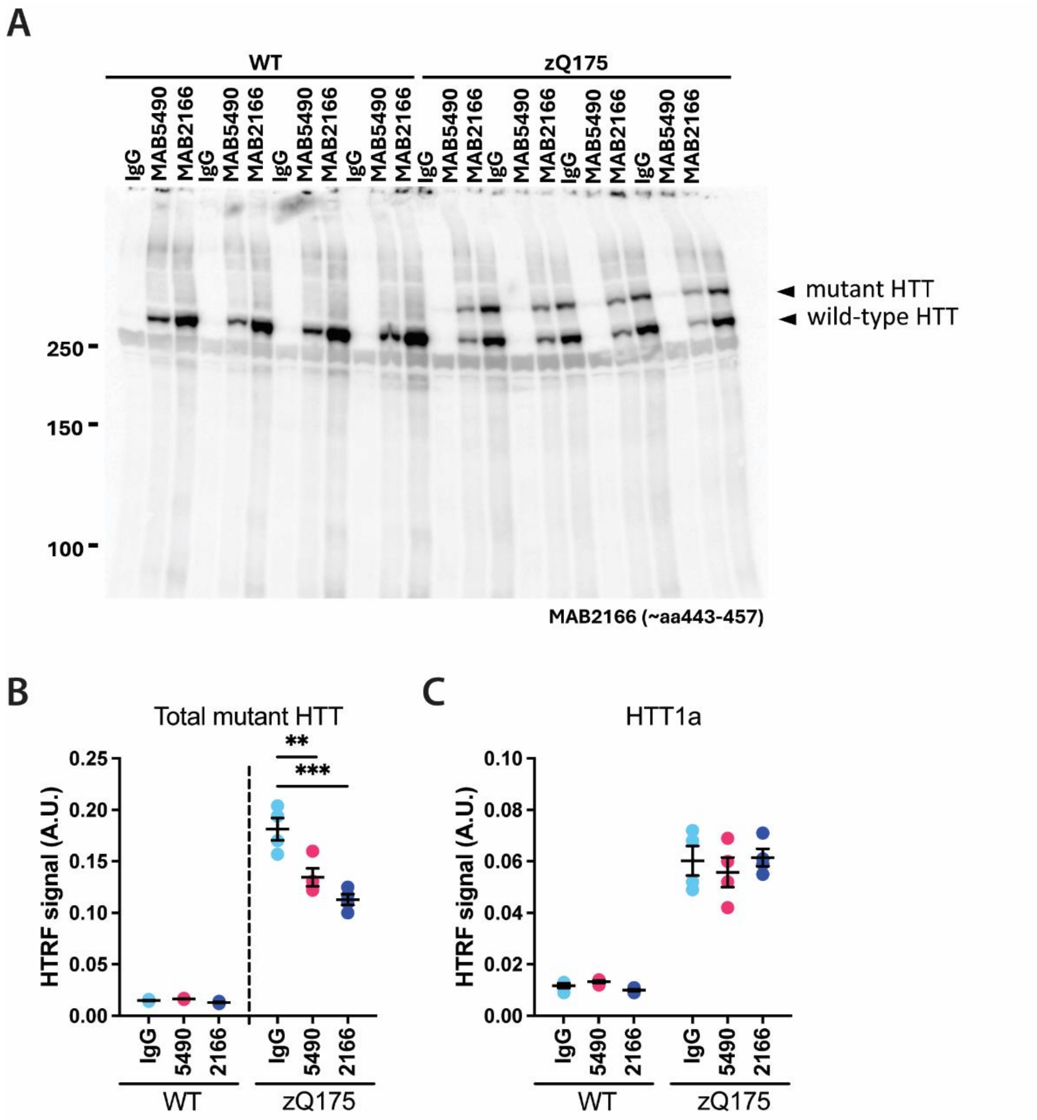


**Fig. S2. Immunodepletion of HTT from cortical lysates does not result in an increase in HTT1a.** (**A**) Immunoprecipitation of mutant HTT with the MAB5490 (amino acids 115-129) or MAB2166 (amino acids 443-457) antibodies from wild-type and zQ175 cortical lysates at 2 months of age and immunodetection with MAB2166. The IgG control was negative. The MAB2166 antibody removed more HTT protein from the lysates than the MAB5490 antibody. Wild-type and mutant HTT were removed from the zQ175 lysates to comparable extents. (n = 4/ immunodepletion/ genotype). Protein standards are in kilodaltons. (**B**) HTRF analysis of the depleted lysates with the 2B7-4C9 antibody pair that detects total mutant HTT confirms that more full-length HTT has been removed from the lysates by immunodepletion with MAB2166. (**C**) HTRF analysis of the depleted lysates with the 2B7-MW8 antibody pair, specific for HTT1a, showed that HTT1a levels were unchanged. Statistical analysis was one-way ANOVA with Tukey’s post hoc correction. Error bars: mean ± SEM. ***$P \leq 0.001$. AU = arbitrary units, WT = wild type.

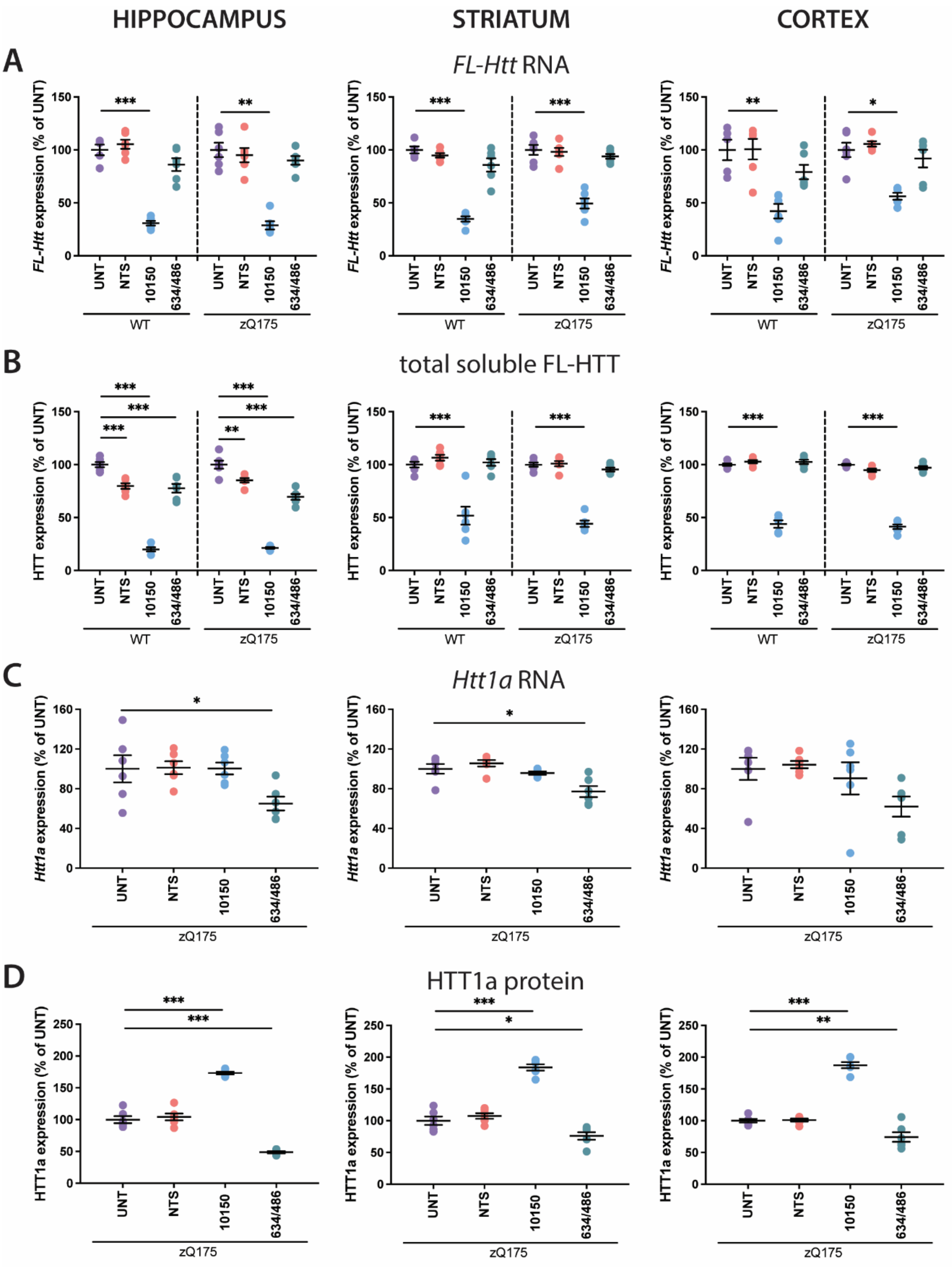


**Fig. S3. Effect of treatment with 10150 or 634/486 on huntingtin mRNA and protein levels one-month post-injection.** Wild-type and zQ175 mice were either untreated or treated with NTS, 10150 or 634/486 at 2 months of age and sacrificed at 3 months. (**A-D**) There was a reduction in full-length HTT levels in NTS treated wild-type and zQ175 mice in the hippocampus, but otherwise treatment with NTS had no effect on full-length *Htt* mRNA or HTT protein, respectively, or the *Htt1a* transcript or HTT1a protein in any brain region. (**A**) Treatment with 10150 resulted in a decrease in full-length *Htt* mRNA in wild-type and zQ175 mice. (**B**) Treatment with 10150 resulted in a decrease in full-length HTT protein

(MAB5490-MAB2166) in wild-type and zQ175 mice. (**C**) Treatment with 634/486 resulted in a decrease in the *Htt1a* transcript in zQ175 mice in the hippocampus and striatum. (**D**) Treatment with 634/486 resulted in a decrease in the HTT1a protein (2B7-MW8) in zQ175 mice in the hippocampus, striatum and cortex. Treatment with 10150 resulted in an apparent increase in HTT1a levels in all three brains regions. (**A-D**) RNA and protein analysis was performed with separate sets of mice n = 3/ gender/ treatment. Statistical analysis was one-way ANOVA with Tukey's post hoc correction or Kruskal-Wallis with Dunn's post hoc correction. Error bars: mean ± SEM. $*P \leq 0.05$, $**P \leq 0.01$, $***P \leq 0.001$. FL = full-length, NTS = non-targeting siRNA, WT = wild type UNT = untreated.

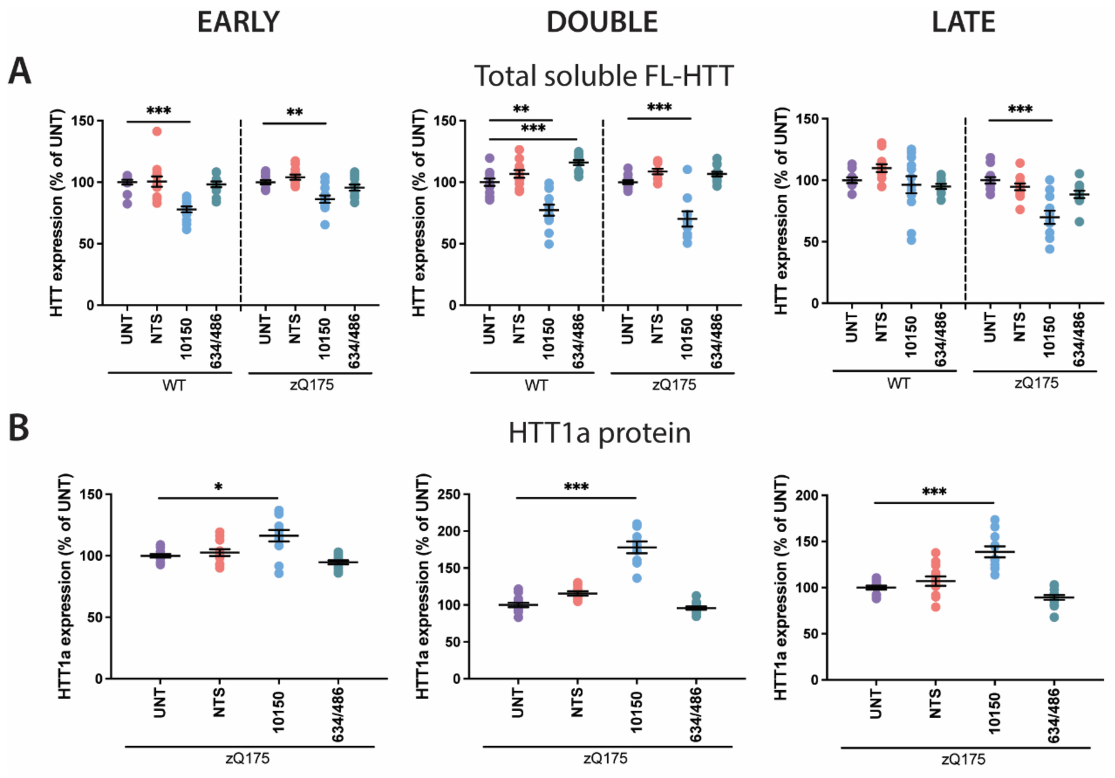


**Fig. S4. Effect of treatment with 10150 or 634/486 on HTT or HTT1a levels in the striatum four months post-injection.** Wild-type and zQ175 mice, either untreated, or 4 months post-treatment with NTS, 10150 or 634/486 in the EARLY, DOUBLE or LATE cohorts. (**A-B**) Treatment with NTS had no effect on full-length HTT or HTT1a proteins in the striatum in any of the treatment cohorts. (**A**) Full-length HTT protein levels (HTRF assay MAB5490-MAB2166) remained decreased in the EARLY (wild-type 22%, zQ175 14%), DOUBLE (wild-type 23%, zQ175 30%) and LATE (wild-type 4%, zQ175 30%) cohorts in the striatum 4 months post-treatment with 10150. There was a large variability in HTT levels between animals, particularly in the LATE cohort. (**B**) There was no reduction in striatal HTT1a protein (HTRF assay 2B7-MW8) in zQ175 mice in any of the cohorts 4 months post-treatment with 634/486. Treatment with 10150 resulted in an apparent increase in HTT1a levels in all three cohorts. n = 5-7 /gender/ treatment. Statistical analysis was one-way ANOVA with Tukey's post hoc correction or Kruskal-Wallis with Dunn's post hoc correction. Error bars: mean ± SEM. *$P \leq 0.05$, **$P \leq 0.01$, ***$P \leq 0.001$. FL = full-length, NTS = non-targeting siRNA, WT = wild type UNT = untreated.

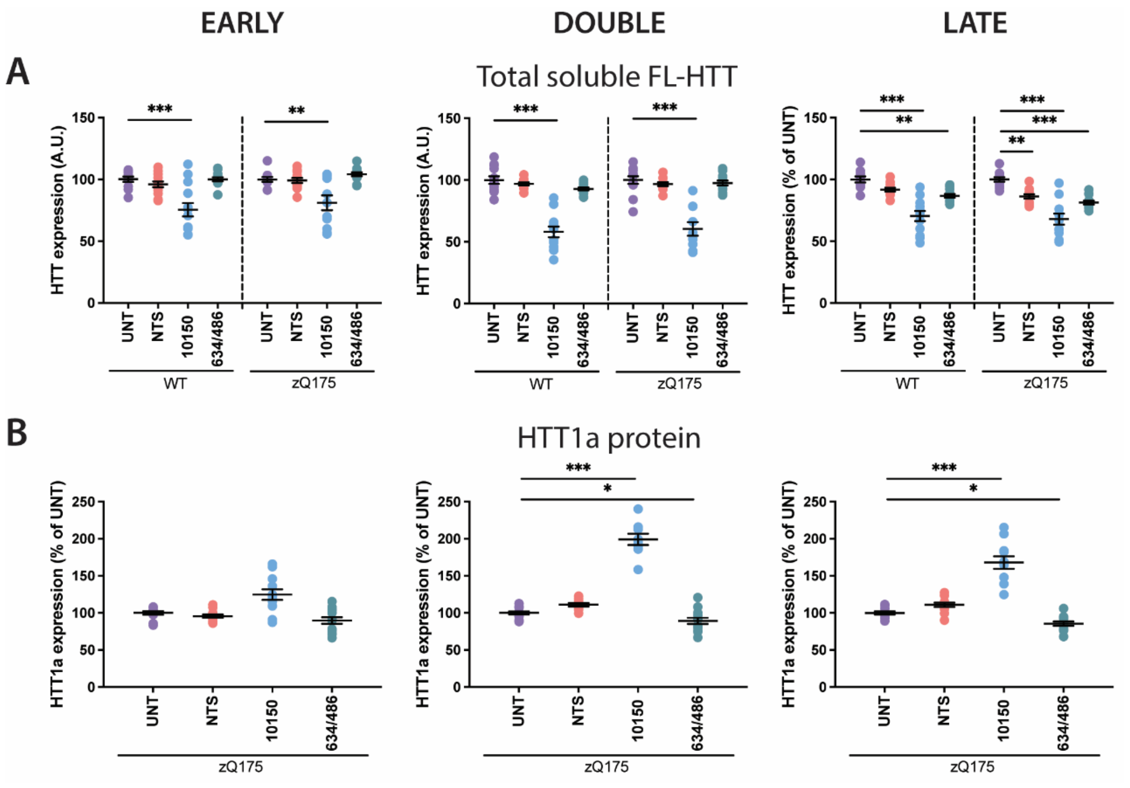


**Fig. S5. Effect of treatment with 10150 or 634/486 on HTT or HTT1a levels in the cortex four months post-injection.** Wild-type and zQ175 mice, either untreated, or 4 months post-treatment with NTS, 10150 or 634/486 in the EARLY, DOUBLE or LATE cohorts. (**A-B**) there was a reduction in full-length HTT protein in NTS treated mice in the LATE cohort only, otherwise, the NTS had no effect on full-length HTT or HTT1a proteins in the cortex. (**A**) Full-length HTT protein levels (HTRF assay MAB5490-MAB2166) remained decreased in the EARLY (wild-type 25%, zQ175 19%), DOUBLE (wild-type 42%, zQ175 40%) and LATE (wild-type 30%, zQ175 32%) cohorts in the cortex 4 months post-treatment with 10150. (**B**) A reduction in cortical HTT1a protein (HTRF assay 2B7-MW8) could be detected in zQ175 mice in the DOUBLE (11%) and LATE (14%) cohorts 4 months post-treatment with 634/486. Treatment with 10150 resulted in an apparent increase in HTT1a levels in all three cohorts, as before. n = 5-7/ gender/ treatment. Statistical analysis was one-way ANOVA with Tukey's post hoc correction or Kruskal-Wallis with Dunn's post hoc correction. Error bars: mean ± SEM. **P* ≤ 0.05, ***P* ≤ 0.01, ****P* ≤ 0.001. FL = full-length, NTS = non-targeting siRNA, WT = wild type UNT = untreated.

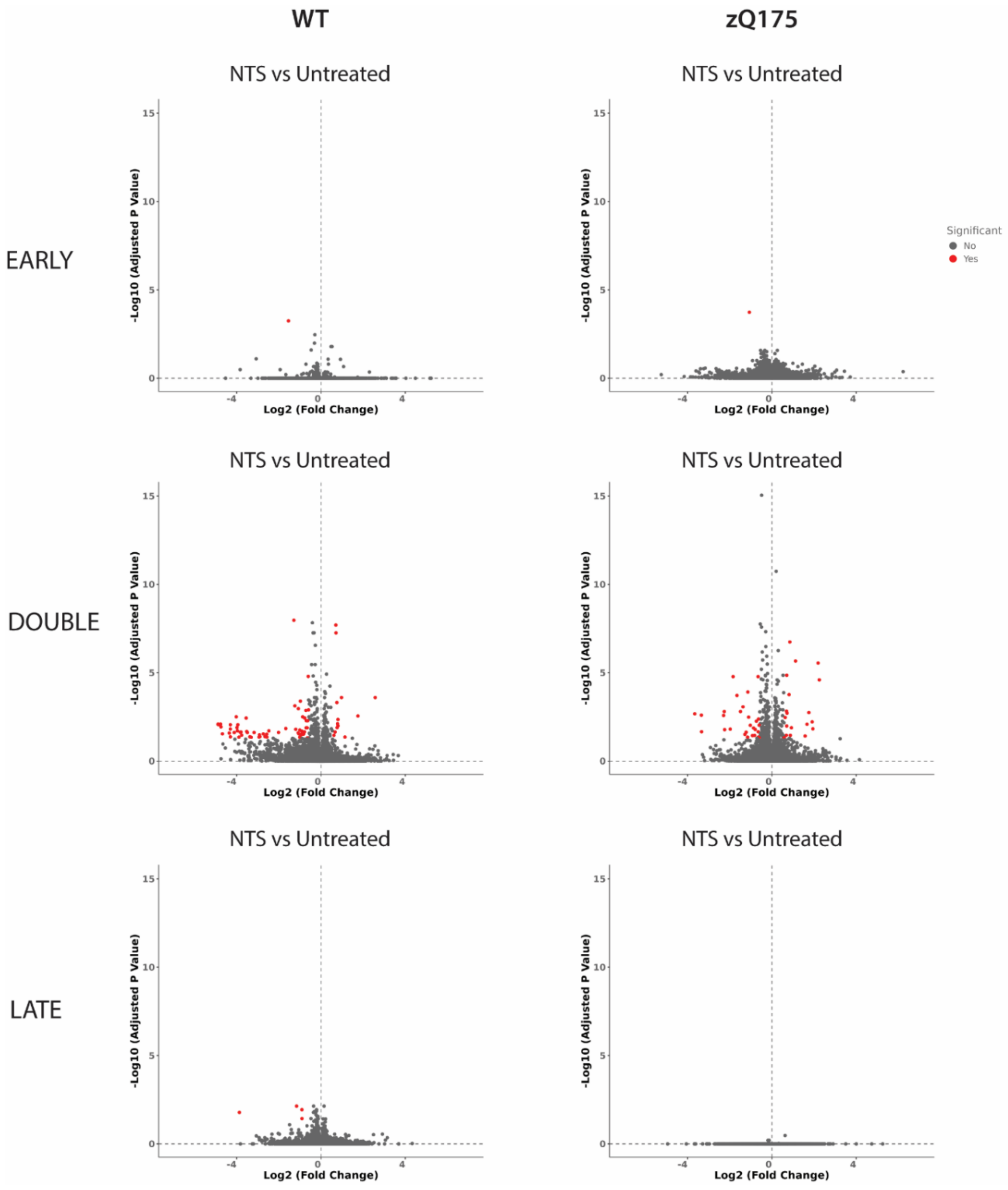


**Fig. S6. Hippocampal transcriptional response to treatment with the non-targeting siRNA.** Volcano plots comparing the transcriptional profile in the hippocampus between NTS and untreated mice for EARLY, DOUBLE and LATE cohorts. Only the wild-type mice and the zQ175 mice for the DOUBLE cohort had a differential transcriptional signature between NTS and untreated animals. The expression of 73 and 47 genes was altered between the wild-type NTS and untreated and zQ175 NTS and untreated groups, respectively, with 11 genes in common. Statistical significance was defined as fold change > 1.5, and adjusted *P*-value < 0.05. NTS = non-targeting siRNA, WT = wild type.

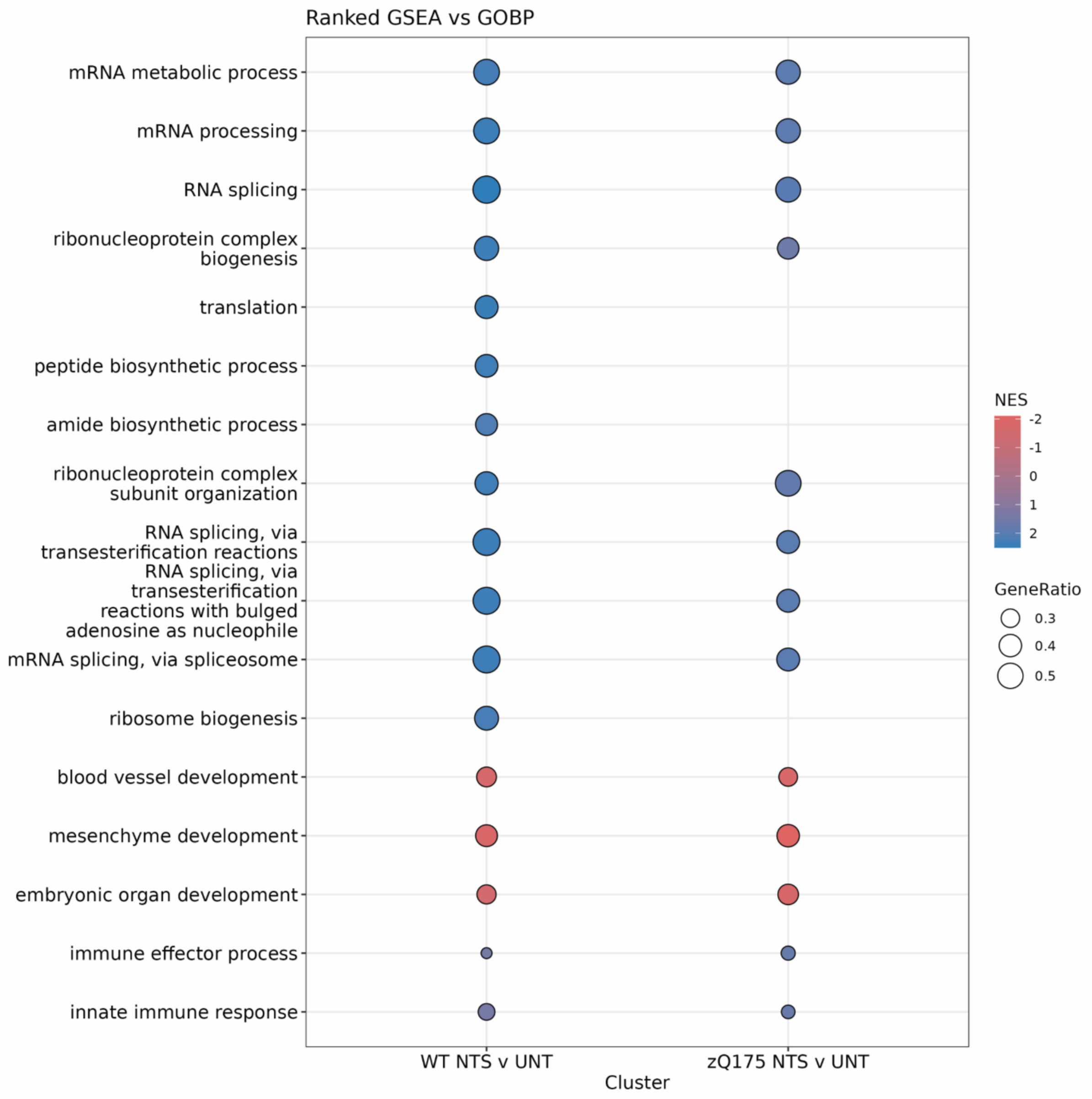


**Fig. S7. Hippocampal transcriptional response to treatment with the non-targeting siRNA in the DOUBLE cohort.** A ranked gene set enrichment analysis vs GOBP for the genes with differential expression levels between NTS and untreated wild-type or zQ175 mice for the DOUBLE cohort. The top-ranking hits are illustrated. Both contrasts show induction of immune response gene sets as well as induction of gene sets related to mRNA processing and splicing. Wild-type mice saw a further induction of gene sets related to amino acid biosynthesis, ribosomes, and translation. GESA = gene set enrichment analysis, GOBP = gene ontology biological process, NES = normalized enrichment score, NTS = non-targeting siRNA, UNT = untreated, WT = wild type.

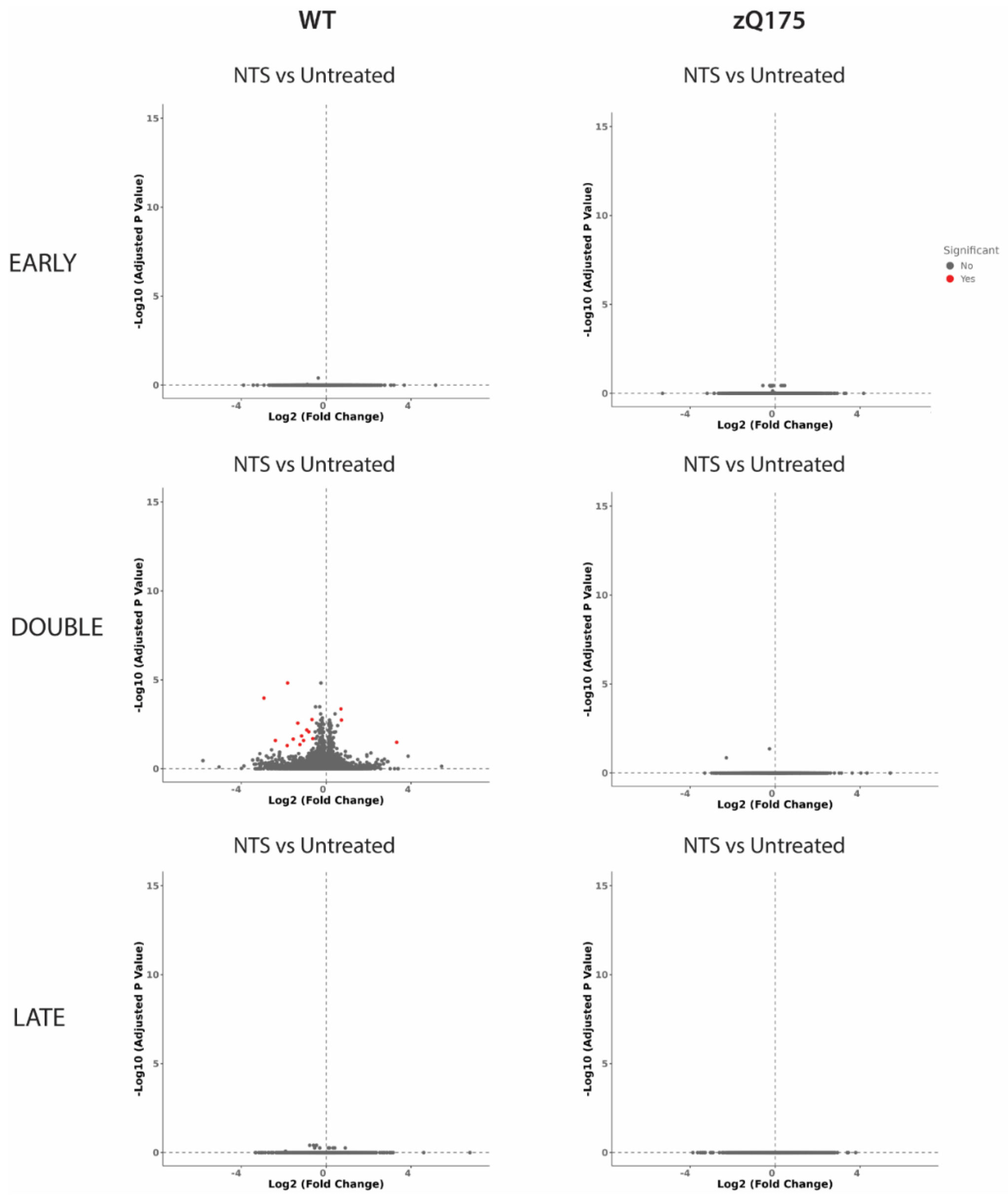


**Fig. S8. Striatal transcriptional response to treatment with the non-targeting siRNA.** Volcano plots comparing the transcriptional profile in the striatum between NTS and untreated mice for EARLY, DOUBLE and LATE cohorts. Only the wild-type mice DOUBLE cohort had a differential transcriptional signature between NTS and untreated animals. This comprised 16 genes that had a statistically significant difference in expression level: *Capn11*, *Grin1os* and *Muc3a* were upregulated and *Aldh1a2*, *Adra1d*, *Col3a1*, *Col18a1*, *Dcn*, *Epn3*, *Igf2*, *Mrc1*, *Prlr*, *Ramp3*, *Slc13a4*, *Srpx2* and 7630403G23Rik were downregulated, numbers likely to reflect the noise in the system. Therefore, treatment with NTS had no effect on the striatal transcriptome. Statistical significance was defined as fold change > 1.5, and adjusted *P*-value < 0.05. NTS = non-targeting siRNA.

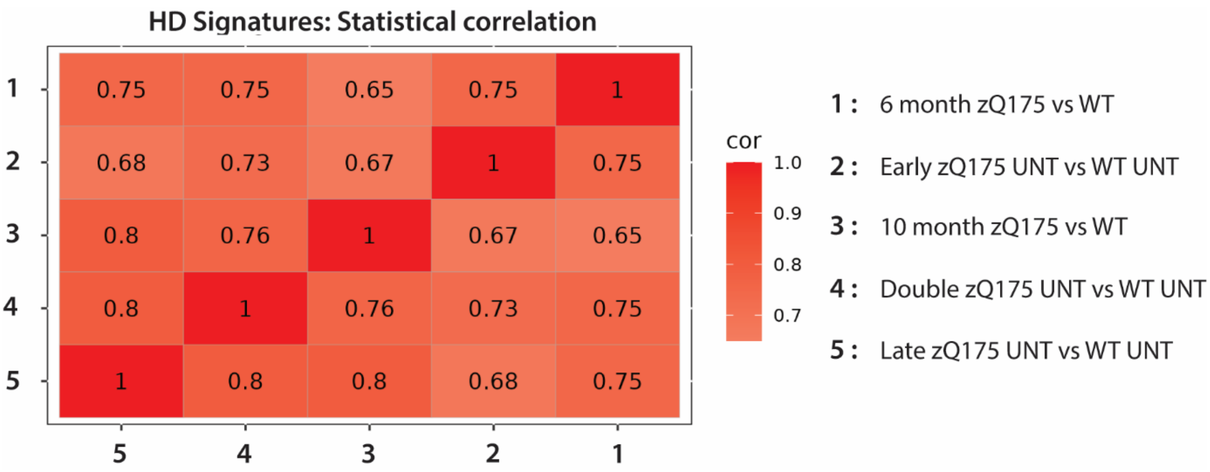


**Fig. S9. Striatal transcriptional profile is comparable to that in previously published data sets**. Heatmap for dysregulated genes in the striatum of untreated of zQ175 and untreated wild-type mice for the EARLY, DOUBLE and LATE cohorts (represented by the DESeq2 Wald statistic) as compared to published data sets zQ175 versus wild-type mice at 6 and 10 months of age from Langfelder et a. (2016). Comparison of the dysregulated genes in these studies are highly correlated. Cor = correlation, M = months, STR = striatum, NTS = non-targeting siRNA, UNT = untreated.

### Table S1. Sequence and modifications of oligonucleotides used in the luciferase screen.

| Name | Strand modifications of *Htt* intron 1 | | Species | | Primary screen |
|---|---|---|---|---|---|
| | Sense strand | Antisense strand | M. musculus position | H. sapiens position | *Htt* mRNA expression % control |
| 316 | (fU)#(mG)#(fC)(mG)(fA)(mA)(fG)(mU)(fU)(mA)(fG)(mG)(fG)#(mA)#(fA)-TegChol | P(mU)#(fU)#(mC)(fC)(mC)(fU)(mA)(fA)(mC)(fU)(mU)(fC)(mG)#(fC)#(mA)#(fA)#(mA)#(fC)#(mU)#(fG) | 604 | N/A | 17.88 |
| 324 | (fU)#(mA)#(fG)(mG)(fG)(mA)(fA)(mC)(fG)(mA)(fA)(mC)(fU)#(mU)#(fA)-TegChol | P(mU)#(fA)#(mA)(fG)(mU)(fU)(mC)(fG)(mU)(fU)(mC)(fC)(mC)#(fU)#(mA)#(fA)#(mC)#(fU)#(mU)#(fC) | 612 | N/A | 18.54 |
| 325 | (fA)#(mG)#(fG)(mG)(fA)(mA)(fC)(mG)(fA)(mA)(fC)(mU)(fU)#(mG)#(fA)-TegChol | P(mU)#(fC)#(mA)(fA)(mG)(fU)(mU)(fC)(mG)(fU)(mU)(fC)(mC)#(fC)#(mU)#(fA)#(mA)#(fC)#(mU)#(fU) | 613 | N/A | 35.18 |
| 342 | (fC)#(mU)#(fC)(mU)(fC)(mU)(fU)(mC)(fU)(mG)(fG)(mA)(fG)#(mA)#(fA)-TegChol | P(mU)#(fU)#(mC)(fU)(mC)(fC)(mA)(fG)(mA)(fA)(mG)(fA)(mG)#(fA)#(mG)#(fA)#(mA)#(fA)#(mC)#(fA) | 630 | N/A | 56.57 |
| 348 | (fU)#(mC)#(fU)(mG)(fG)(mA)(fG)(mA)(fA)(mA)(fC)(mU)(fG)#(mG)#(fA)-TegChol | P(mU)#(fC)#(mC)(fA)(mG)(fU)(mU)(fU)(mC)(fU)(mC)(fC)(mA)#(fG)#(mA)#(fA)#(mG)#(fA)#(mG)#(fA) | 636 | N/A | 68.79 |
| 385 | (fU)#(mG)#(fA)(mA)(fG)(mA)(fG)(mA)(fA)(mC)(fU)(mU)(fG)#(mG)#(fA)-TegChol | P(mU)#(fC)#(mC)(fA)(mA)(fG)(mU)(fU)(mC)(fU)(mC)(fU)(mU)#(fC)#(mA)#(fC)#(mA)#(fA)#(mC)#(fA) | 673 | N/A | 28.85 |
| 416 | (fG)#(mG)#(fG)(mU)(fU)(mA)(fC)(mC)(fU)(mC)(fC)(mU)(fC)#(mA)#(fA)-TegChol | P(mU)#(fU)#(mG)(fA)(mG)(fG)(mA)(fG)(mG)(fU)(mA)(fA)(mC)#(fC)#(mC)#(fU)#(mA)#(fG)#(mA)#(fG) | 704 | N/A | 25.63 |
| 477 | (fU)#(mA)#(fG)(mU)(fG)(mG)(fA)(mU)(fG)(mA)(fC)(mA)(fU)#(mA)#(fA)-TegChol | P(mU)#(fU)#(mA)(fU)(mG)(fU)(mC)(fA)(mU)(fC)(mC)(fA)(mC)#(fU)#(mA)#(fC)#(mC)#(fC)#(mG)#(fC) | 765 | N/A | 25.47 |
| 478 | (fA)#(mG)#(fU)(mG)(fG)(mA)(fU)(mG)(fA)(mC)(fA)(mU)(fA)#(mA)#(fA)-TegChol | P(mU)#(fU)#(mU)(fA)(mU)(fG)(mU)(fC)(mA)(fU)(mC)(fC)(mA)#(fC)#(mU)#(fA)#(mC)#(fC)#(mC)#(fG) | 766 | N/A | 20.43 |
| 486 | (fA)#(mC)#(fA)(mU)(fA)(mA)(fU)(mG)(fC)(mU)(fU)(mU)(fU)#(mA)#(fA)-TegChol | P(mU)#(fU)#(mA)(fA)(mA)(fA)(mG)(fC)(mA)(fU)(mU)(fA)(mU)#(fG)#(mU)#(fC)#(mA)#(fU)#(mC)#(fC) | 774 | 515 | 11.61 |
| 538 | (fA)#(mA)#(fC)(mG)(fC)(mA)(fU)(mC)(fC)(mA)(fA)(mU)(fG)#(mG)#(fA)-TegChol | P(mU)#(fC)#(mC)(fA)(mU)(fU)(mG)(fG)(mA)(fU)(mG)(fC)(mG)#(fU)#(mU)#(fC)#(mA)#(fC)#(mA)#(fC) | 826 | N/A | 81.80 |
| 577 | (fG)#(mA)#(fA)(mG)(fC)(mA)(fG)(mC)(fC)(mU)(fG)(mU)(fG)#(mA)#(fA)-TegChol | P(mU)#(fU)#(mC)(fA)(mC)(fA)(mG)(fG)(mC)(fU)(mG)(fC)(mU)#(fU)#(mC)#(fA)#(mA)#(fG)#(mU)#(fG) | 865 | N/A | 51.42 |
| 614 | (fG)#(mG)#(fC)(mG)(fU)(mU)(fU)(mC)(fA)(mU)(fU)(mU)(fA)#(mG)#(fA)-TegChol | P(mU)#(fC)#(mU)(fA)(mA)(fA)(mU)(fG)(mA)(fA)(mA)(fC)(mG)#(fC)#(mC)#(fA)#(mG)#(fG)#(mA)#(fG) | 902 | N/A | 40.07 |
| 619 | (fU)#(mU)#(fC)(mA)(fU)(mU)(fU)(mA)(fG)(mU)(fU)(mU)(fG)#(mU)#(fA)-TegChol | P(mU)#(fA)#(mC)(fA)(mA)(fA)(mC)(fU)(mA)(fA)(mA)(fU)(mG)#(fA)#(mA)#(fA)#(mC)#(fG)#(mC)#(fC) | 907 | N/A | 24.38 |
| 631 | (fG)#(mU)#(fG)(mG)(fU)(mG)(fU)(mA)(fG)(mU)(fG)(mU)(fA)#(mG)#(fA)-TegChol | P(mU)#(fC)#(mU)(fA)(mC)(fA)(mC)(fU)(mA)(fC)(mA)(fC)(mC)#(fA)#(mC)#(fA)#(mA)#(fA)#(mC)#(fU) | 919 | N/A | 7.04 |

| | | | | | |
|---|---|---|---|---|---|
| 634 | (fG)#(mU)#(fG)(mU)(fA)(mG)(fU)(mG)(fU)(mA)(fG)(mU)(fU) #(mA)#(fA)-TegChol | P(mU)#(fU)#(mA)(fA)(mC)(fU)(mA)(fC)(mA)(fC)(mU)(fA)(mC)#(fA)# (mC)#(fC)#(mA)#(fC)#(mA)#(fA) | 922 | N/A | 8.27 |
| 635 | (fU)#(mG)#(fU)(mA)(fG)(mU)(fG)(mU)(fA)(mG)(fU)(mU)(fA) #(mA)#(fA)-TegChol | P(mU)#(fU)#(mU)(fA)(mA)(fC)(mU)(fA)(mC)(fA)(mC)(fU)(mA)#(fC)# (mA)#(fC)#(mC)#(fA)#(mC)#(fA) | 923 | N/A | 14.13 |
| 636 | (fG)#(mU)#(fA)(mG)(fU)(mG)(fU)(mA)(fG)(mU)(fU)(mA)(fA) #(mA)#(fA)-TegChol | P(mU)#(fU)#(mU)(fU)(mA)(fA)(mC)(fU)(mA)(fC)(mA)(fC)(mU)#(fA) #(mC)#(fA)#(mC)#(fC)#(mA)#(fC) | 924 | N/A | 22.64 |
| 643 | (fA)#(mG)#(fU)(mU)(fA)(mA)(fA)(mC)(fC)(mA)(fG)(mG)(fU)# (mU)#(fA)-TegChol | P(mU)#(fA)#(mA)(fC)(mC)(fU)(mG)(fG)(mU)(fU)(mU)(fA)(mA)#(fC) #(mU)#(fA)#(mC)#(fA)#(mC)#(fU) | 931 | N/A | 15.16 |
| 664 | (fU)#(mA)#(fA)(mA)(fC)(mC)(fA)(mG)(fG)(mU)(fU)(mU)(fU)# (mA)#(fA)-TegChol | P(mU)#(fU)#(mA)(fA)(mA)(fA)(mC)(fC)(mU)(fG)(mG)(fU)(mU)#(fU) #(mA)#(fA)#(mC)#(fU)#(mA)#(fC) | 934 | N/A | 61.43 |
| 646 | (fA)#(mA)#(fA)(mC)(fC)(mA)(fG)(mG)(fU)(mU)(fU)(mU)(fA)# (mA)#(fA)-TegChol | P(mU)#(fU)#(mU)(fA)(mA)(fA)(mA)(fC)(mC)(fU)(mG)(fG)(mU)#(fU) #(mU)#(fA)#(mA)#(fC)#(mU)#(fA) | 935 | N/A | 25.94 |
| 647 | (fU)#(mA)#(fG)(mC)(fC)(mA)(fG)(mA)(fG)(mA)(fG)(mG)(fU)# (mG)#(fA)-TegChol | P(mU)#(fC)#(mA)(fC)(mC)(fU)(mC)(fU)(mC)(fU)(mG)(fG)(mC)#(fU)# (mA)#(fU)#(mG)#(fC)#(mU)#(fU) | 952 | N/A | 19.35 |
| 696 | (fG)#(mG)#(fC)(mA)(fG)(mU)(fU)(mG)(fG)(mA)(fU)(mG)(fA)# (mG)#(fA)-TegChol | P(mU)#(fC)#(mU)(fC)(mA)(fU)(mC)(fC)(mA)(fA)(mC)(fU)(mG)#(fC)# (mC)#(fU)#(mG)#(fC)#(mA)#(fG) | 984 | N/A | 37.93 |
| 713 | (fU)#(mA)#(fU)(mU)(fU)(mG)(fU)(mC)(fA)(mA)(fG)(mU)(fA)# (mC)#(fA)-TegChol | P(mU)#(fG)#(mU)(fA)(mC)(fU)(mU)(fG)(mA)(fC)(mA)(fA)(mA)#(fU) #(mA)#(fC)#(mA)#(fA)#(mC)#(fU) | 1001 | N/A | 7.98 |

Detailed sequence, chemical modification patterns, and efficacy of siRNAs. Mouse huntingtin accession number - NC_000071.7. Chemical modifications are designated as follows. “#” –phosphorothioate bond, “m” – 2’-O-Methyl, “f” – 2’-Fluoro, “P” – 5’ Phosphate, “tegChol” – tetraethylene glycol (teg)-cholesterol. All sequences are homologous to mouse huntingtin.

### Table S2. Sequence and modifications of oligonucleotides used for the *in vivo* experiments.

| | | | Strand modifications of *Htt* intron 1 | |
|---|---|---|---|---|
| Gene | Name | Position | Sense strand | Antisense strand |
| *Htt* intron 1 | 486 | 774 | (fA)#(mC)#(fA)(mU)(fA)(mA)(fU)(mG)(fC)(mU)(fU)(mU)(fU)#(mA)#(fA)-DIO | V(mU)#(fU)#(mA)(fA)(mA)(fA)(mG)(fC)(mA)(fU)(mU)(fA)(mU)#(fG)#(mU)#(fC)#(mA)#(fU)#(mC)#(fC) |
| *Htt* intron 1 | 634 | 922 | (fG)#(mU)#(fG)(mU)(fA)(mG)(fU)(mG)(fU)(mA)(fG)(mU)(fU)#(mA)#(fA)-DIO | V(mU)#(fU)#(mA)(fA)(mC)(fU)(mA)(fC)(mA)(fC)(mU)(fA)(mC)#(fA)#(mC)#(fC)#(mA)#(fC)#(mA)#(fA) |
| *Human HTT* | 10150 | 10150 | (fC)#(mA)#(fG)(mU)(fA)(mA)(fA)(mG)(fA)(mG)(fA)(mU)(fU)#(mA)#(fA)-DIO | V(mU)#(fU)#(mA)(fA)(mU)(fC)(mU)(fC)(mU)(fU)(mU)(fA)(mC)#(fU)#(mG)#(fA)#(mU)#(fA)#(mU)#(fA) |
| Non-targeting | NTS | N/A | (fU)#(mG)#(fA)(mC)(fA)(mA)(fA)(mU)(fA)(mC)(fG)(mA)(fU)#(mU)#(mA)-DIO | V(mU)#(fA)#(mA)(fU)(mC)(fG)(mU)(fA)(mU)(fU)(mU)(fG)(mU)#(fC)#(mA)#(fA)#(mU)#(fC)#(mA)#(fU) |

Detailed sequence, chemical modification patterns, and efficacy of siRNAs. Human huntingtin accession number - NM_002111 .6, mouse huntingtin accession number - NC_000071.7. Chemical modifications are designated as follows. "#" –phosphorothioate bond, "m" – 2'-O-Methyl, "f" – 2'-Fluoro, "V" – 5' vinyl-phosphonate, "DIO" – divalent. 486 and 634 homologous to mouse huntingtin, 10150 homologous to mouse and human huntingtin.

**Table S3. CAG repeat length for each of the treatment groups.**

| | Cohort | | | |
|---|---|---|---|---|
| Treatment group | BASELINE | EARLY | DOUBLE | LATE |
| Untreated | 190.7 ± 3.54 | 191.8 ± 4.78 | 190.8 ± 4.38 | 190.6 ± 3.18 |
| NTS | 191.5 ± 2.39 | 190.8 ± 4.23 | 191.7 ± 4.89 | 190.2 ± 5.08 |
| 10150 | 189.7 ± 3.76 | 191.0 ± 3.74 | 191.3 ± 4.37 | 190.0 ± 4.12 |
| 486/634 | 192.1 ± 3.90 | 191.0 ± 4.20 | 191.2 ± 4.46 | 191.0 ± 4.84 |

NTS = non-targeting siRNA. Mean ± Standard deviation.

**Table S4. Antibodies.**

| Name | Immunogen | Epitope | Species | Reference / Source |
|---|---|---|---|---|
| 2B7 | Human HTT peptide: aa 1-17 (*49*) | LMKAFE* | Mouse Monoclonal | CHDI Foundation |
| 3B5H10 | GST-HTT N171 (66Q) (*50*) | PolyQ | Mouse monoclonal | Sigma-Aldrich P1874 |
| 4C9 | Human HTT peptide: aa 51-71 (*51*) | | Mouse Monoclonal | CHDI Foundation |
| MW8 | AEEPLHRPK (67Q) (*52*) | Within: AEEPLHRP (*53*) Ends at proline | Mouse Monoclonal | CHDI Foundation |
| S830 | Exon 1 HTT (53Q) (*54*) | N/A | Sheep polyclonal | In-house |
| MAB5490 | Human HTT: aa 115-129 (*55*) | Within: QSVRNSPEFQKLLGI (mouse L) | Mouse Monoclonal | Sigma-Aldrich, MAB5490 |
| MAB2166 | HTT fusion protein: aa 181-810 (*56*) | Within aa 443-457 GKVLLGEEEALEDDS (*57*) | Mouse Monoclonal | Sigma-Aldrich, MAB2166 |

*Information provided by the CHDI Foundation

**Table S5. Antibody and lysate concentrations for HTRF assays.**

| Antibody Pairing | HTT isoform | Donor (ng / well) | Acceptor (ng / well) | Lysate Concentration |
|---|---|---|---|---|
| MAB5490-Tb: MAB2166-d2 | Full-length HTT | 1 ng | 20 ng | 5% |
| 2B7-Tb: MW8-d2 | HTT1a | 1 ng | 40 ng | 10%<br>5% - immunodepletion |
| 2B7-Tb: 4C9-488 | Mutant HTT | 1 ng | 10 ng | 5% |
| 4C9-Tb: MW8-d2 | HTT aggregation | 1 ng | 10 ng | 10% Crude Lysate |
